# Chemogenetic Hydrogen Sulfide Rescues Vascular Senescence in Ageing

**DOI:** 10.64898/2026.09.29.755422

**Authors:** Khatereh Shabanian, Taraneh Shabanian, Lorenzo Da Dalt, Manuel Colucci, Emrah Eroglu, Hector Rodriguez Cetina Biefer, Omer Dzemali, Matthias Hermann, Andrea Alimonti, Giuseppe Danilo Norata, Francesco Paneni, Frank Ruschitzka, Seyed Soheil Saeedi Saravi

## Abstract

Vascular senescence is a central driver of whole-body ageing, yet rescue strategies remain limited. Here we show that hydrogen sulfide (H_₂_S) bioavailability declines in ageing and atherosclerosis and is associated with endothelial senescence. Developing a novel remote-control, endothelial-specific, mitochondria-targeted chemogenetic system, we spatiotemporally controlled H_₂_S generation and thereby mitigated endothelial senescence-driven programs and reversed vascular remodeling in aged mice.

## Introduction

Vascular senescence acts as both early sensor and driver of whole-body ageing^1,2^. Accumulation of senescent endothelial cells promotes cardiovascular disease, thereby accelerating ageing process^3,4^. While the mechanisms that govern vascular senescence and strategies capable of rescuing this process remain to be fully uncovered, hydrogen sulfide (H_2_S) has been emerged as a critical endogenous modulator of vascular homeostasis^5,6^ and healthy lifespan^7^. Mammals essentially need H_2_S for post-translational protein persulfidation to regulate redox signalling and mitochondrial bioenergetics and improve stress resistance^7^. Experimental models exhibit that elevating H_2_S by endogenous production or donors exerts anti-aging effects and expands lifespan^8^. Mitochondrial H_2_S can also improve mitochondrial dysfunction, a hallmark of ageing^9^. Yet, currently available H_2_S-releasing compounds, including sulfide salts e.g. sodium hydrosulfide (NaHS) and slow-releasing donors, cannot introduce controlled and consistent H_2_S levels, exert off-target effects, and widely fail to mimic endogenous H_2_S signaling spatiotemporally in specific cell types^10^. Varying and non-specific H_2_S production may lead to cytotoxicity or long-term insufficiency. Under physiological conditions in mammals, H_2_S is primarily generated from enzymatic L-cysteine (L-Cys) conversion by cystathionine β-synthase (CBS) and cystathionine γ-lyase (CGL, also known as CTH)^11^. Alternatively, D-cysteine (D-Cys) can be metabolized by a pathway involving d-amino acid oxidase (DAAO) and β-mercaptopyruvate sulfurtransferase (3-MPST)^12,13^. Notably, the canonical H_2_S-generating enzymes decline with ageing^9^, and DAAO is only primarily expressed in peroxisomes of kidney cells and hepatocytes, but not in vascular cells^14^. These suggest that impaired sulfide signaling may contribute to age-associated vascular decline^15^. However, whether loss of endogenous H_2_S represents a modifiable driver of vascular senescence remains unknown.

## Results and Discussion

To address this question, we first examined circulating H_₂_S levels in a human ageing cohort comprising 86 healthy individuals across age groups. We found that plasma H_2_S levels decline with age (aged *vs.* young: median [IQR]: 0.092 [0.054–0.155] µM *vs.* 0.287 [0.157– 0.371] µM; Fig. 1a). Similarly, aged mice also exhibited a significant reduction of H_2_S levels in aortic tissues (Fig. 1b). Beyond H_2_S level reduction, aged mice displayed loss of its enzymatic source in aortic endothelial cells (AECs), evidenced by reduced expression of CTH protein (Fig. 1c). Confirming the crucial functional role of H_2_S in vasculature^16,17^, we revealed that plasma H_2_S levels positively correlated with endothelium-dependent maximal vasorelaxation responses (*R*=0.5682, *P*<0.0001) (Fig. 1d). These emphasize animals with higher H_2_S display stronger vasodilatory responses, supporting a protective role for H_2_S signaling in vascular physiology. Given that endothelial senescence is a major driver of age-associated vascular dysfunction^2^, we next investigated whether H_2_S deficiency was associated with senescence phenotype in AECs. The observations were corroborated by senescence phenotype in AECs, evidenced by marked upregulation of CDK inhibitors *Cdkn2a* (p16^INK4a^), *Cdkn2d* (p19^INK4d^), and *Cdkn1a* (p21^WAF1/Cip1^), and the senescence-associated secretory phenotype (SASP) components *Il1b*, *Il6*, and *Ccl2*, at mRNA level (Fig. 1e).

**Fig. 1.**
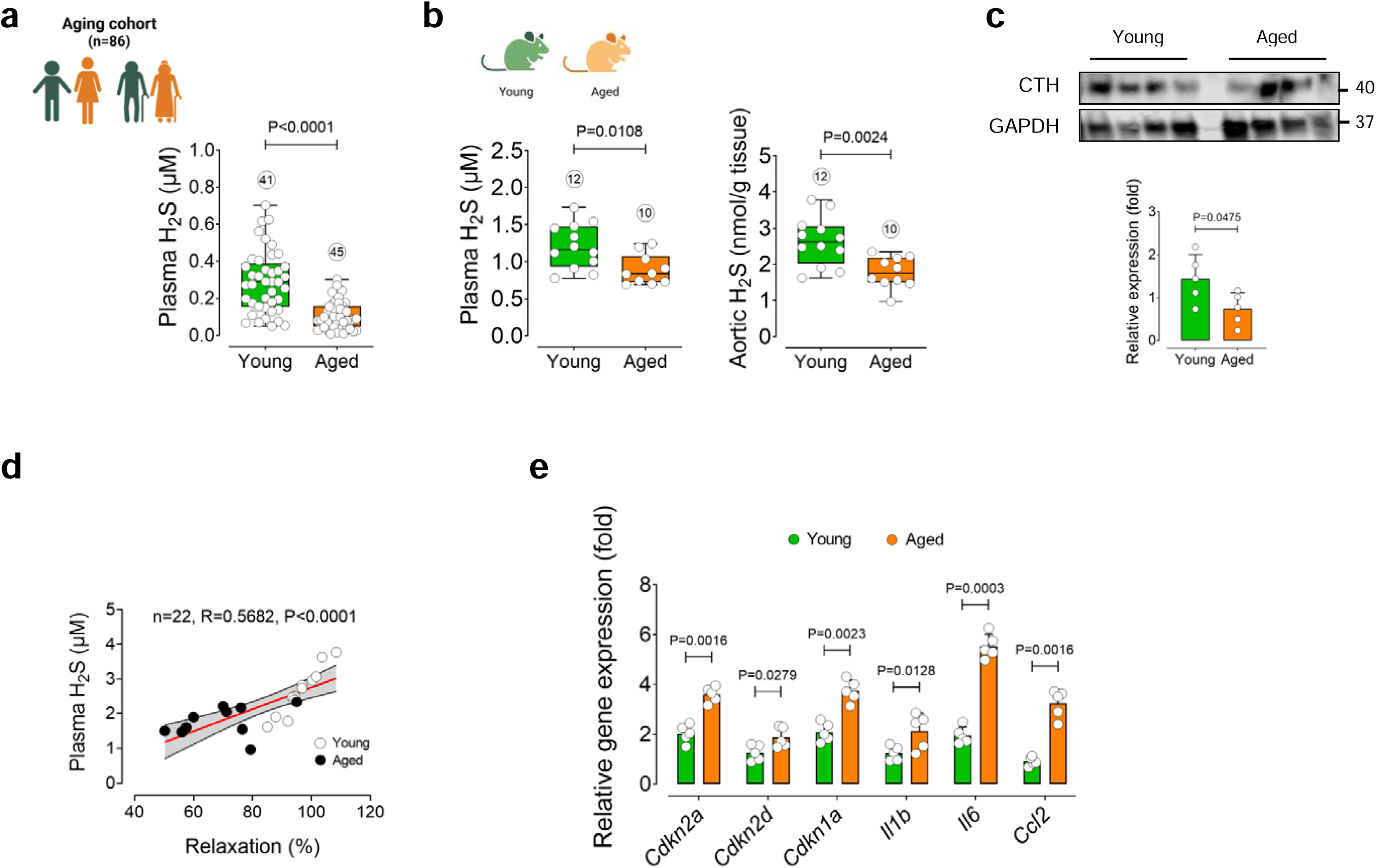
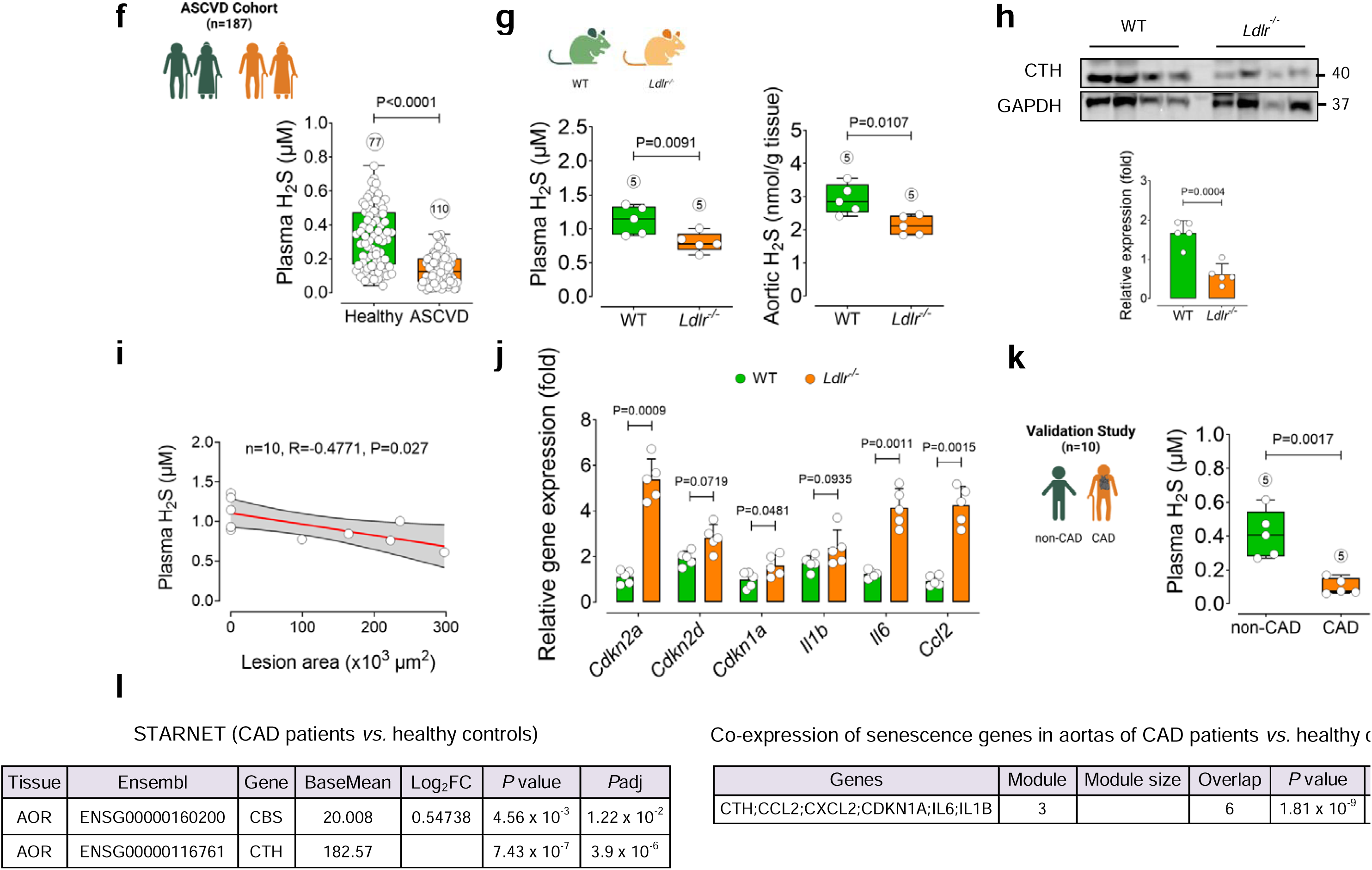
H_2_S declines with age and vascular pathologies. **a,** Plasma H_2_S concentration in the healthy young (18-40 years) and old (>65 years) individuals from the ageing cohort (young, *n*=41; old, *n*=45). **b,** H_2_S concentrations in the plasma and aortic tissues from young (3-month-old) and aged (>24-month-old) mice (young, *n*=12; old, *n*=10). **c,** Representative immunoblots and quantification of intensities for CTH in aortic ECs from young and aged mice (*n*=5). **d,** Correlation between plasma H_2_S levels and maximal vasorelaxation responses (%) to acetylcholine at M in the mouse ageing cohort (*n*=10-12). **e,** qPCR shows transcriptional alterations of CDK inhibitors *Cdkn2a*, *Cdkn2d*, and *Cdkn1a* and SASP components *Ilb*, *Il6*, and *Ccl2* in aortic ECs from young and aged mice (*n*=5). **f,** Plasma H_2_S concentration in aged ASCVD patients (>80 years old) enrolled in the ASCVD cohort (*n*=110; male and female) and healthy controls (*n*=77; male and female). **g,h,** H_2_S concentrations in the plasma and aortic tissues **(g)** and representative immunoblots and quantification of intensities for CTH in aortic ECs **(h)** from WT mice fed a chow and *Ldlr^−/−^* mice fed a Western diet for 12 weeks (*n*=5). **i,** Correlation of plasma H_2_S levels with aortic root lesion area (%). **j,** qPCR shows transcriptional alterations of CDK inhibitors *Cdkn2a*, *Cdkn2d*, and *Cdkn1a* and SASP components *Ilb*, *Il6*, and *Ccl2* in aortic ECs from WT and *Ldlr^−/−^* mice (*n*=5). **k,** Plasma H_2_S levels in aged CAD patients undergoing CABG surgery (*n*=5) and non-CAD controls (*n*=5) in the validation study. **l,** Expression of *CBS* and *CTH* genes (upper) and co-expression of *CTH* with the senescence-associated genes *CDKN1A*, *IL1B*, *IL6*, *CCL2*, and *CXCL2* in atherosclerotic aortic wall (AOR) of patients with CAD (*n*=600) and healthy individuals (*n*=250) in the STARNET database. Error bars represent SD (**a,b,c,e,f,g,h,j,k**). Statistical analysis was performed using two-tailed unpaired Student’s *t*-test (**a,b,c,e,f,g,h,j,k**) and two-sided Spearman’s rank correlation test (**D,I**). Data are shown as median with min–max; each box-plot represents interquartile range (IQR); center lines indicate the median; box-plots extend from the min to max values (**a,b,f,g,k**). Images created with BioRender.com (**a,b,f,g,k**).

Atherosclerosis represents a prototypical disease of accelerated vascular ageing^18,19^. We, therefore, tested whether similar H_₂_S deficiency is also associated with human atherosclerotic cardiovascular disease (ASCVD). Plasma H_2_S concentrations were quantified in the ASCVD cohort (Aging Heart Zurich Cohort), comprising 110 aged ASCVD patients (>80 years old) and 77 non-ASCVD controls. As shown in Fig. 1f, circulating H_2_S levels were significantly lower in ASCVD patients compared to control counterparts (median [IQR]: 0.121 [0.063–0.205] µM *vs.* 0.332 [0.167–0.472] µM). Consistent with these clinical observations, atherosclerosis-prone *Ldlr^−/-^* mice fed a Western diet (WD; high-cholesterol diet) exhibited significantly lower H_2_S levels compared to wild-type mice maintained on chow diet (Fig. 1g). Immunoblots confirmed pronounced downregulation of endothelial CTH in WD-fed *Ldlr^−/-^* mice (Fig. 1h). Importantly, H_2_S concentrations inversely correlated with aortic root lesion area (*R*=0.4771, *P*=0.027; Fig. 1i; Supplemental Fig. 1), reinforcing a robust association between H_2_S and atherosclerosis burden. These were accompanied by a robust activation of endothelial senescence programs, including upregulation of CDK inhibitors *Cdkn2a* and *Cdkn1a*, and SASP components *Il6* and *Ccl2* (Fig. 1j). As a proof of concept in humans, reduced H_2_S levels were observed in aged patients with coronary artery disease (CAD) undergoing CABG surgery compared with non-CAD controls in a validation study (Fig. 1k). To extend these findings, we analyzed vascular transcriptomic datasets from the Stockholm-Tartu Atherosclerosis Reverse Network Engineering Task (STARNET) database^20^. Data showed significantly reduced expression of H_2_S–producing enzymes, *CBS* and *CTH*, alongside the co-expression network with senescence-related genes, including *CDKN1A*, *IL1B*, *IL6*, *CCL2*, and *CXCL2*, in atherosclerotic aortic tissues from CAD patients compared with healthy controls (Fig. 1l). Together, these findings support a conserved crosstalk between H_2_S deficiency, endothelial senescence-driven program and atherosclerotic CVD.

We next developed a tractable chemogenetic system to restore H_2_S signaling and reverse endothelial senescence. To enable precise spatiotemporal control of H_2_S production, we employed a chemogenetic tool that permits inducible generation of H_2_S in cell mitochondria. This system utilizes a recombinant yeast D-amino acid oxidase (DAAO), which stereoselectively oxidizes D-amino acids but remains inactive in mammalian cells that primarily contain L-amino acids^21,22^. Upon administration of exogenous D-Cys, DAAO converts the substrate into β-mercaptopyruvate (3MP) that is subsequently converted to H_2_S and pyruvate by mitochondrial 3-MPST (Fig. 2a). Therefore, this tool allows spatiotemporally switching on-or-off chemogenetic H_2_S production. To monitor mitochondrial H_2_S dynamics, we engineered a mitochondria-targeted fusion construct combining DAAO with a genetically encoded H_₂_S biosensor hsGFP. This design allows simultaneous production and detection of H_₂_S in the same subcellular compartment. Replicative senescent human aortic endothelial cells (HAEC; passages 15–17) were transduced with a lentiviral construct encoding mitochondria-targeted DAAO-hsGFP and further exposed to either D-Cys or L-Cys (as control). Real-time fluorescence imaging revealed a rapid and robust increase in mitochondrial H_₂_S levels exclusively in D-Cys-treated cells, confirming the functionality of our chemogenetic system (Fig. 2b,c). Notably, D-Cys exerts no cytotoxic effects at the intended concentration (Supplemental Fig. 2). As a proof of concept for our successful D-Cys-mediated mitochondrial H_2_S generation, we also tested another chemogenetic tool containing D-cysteine desulfhydrase (stDCyD)—recently established for controlled H_2_S manipulation^12^.

**Fig. 2.**
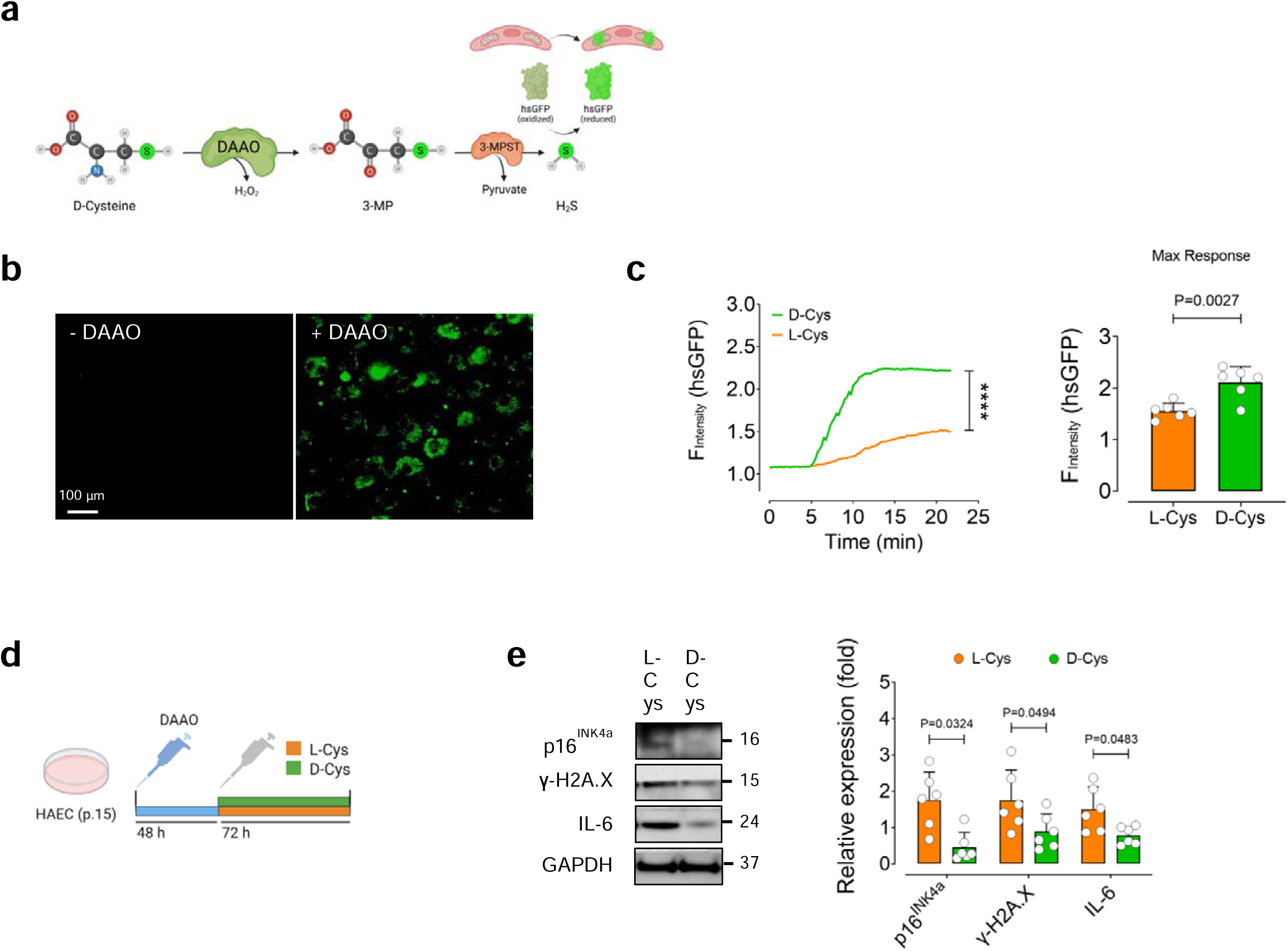
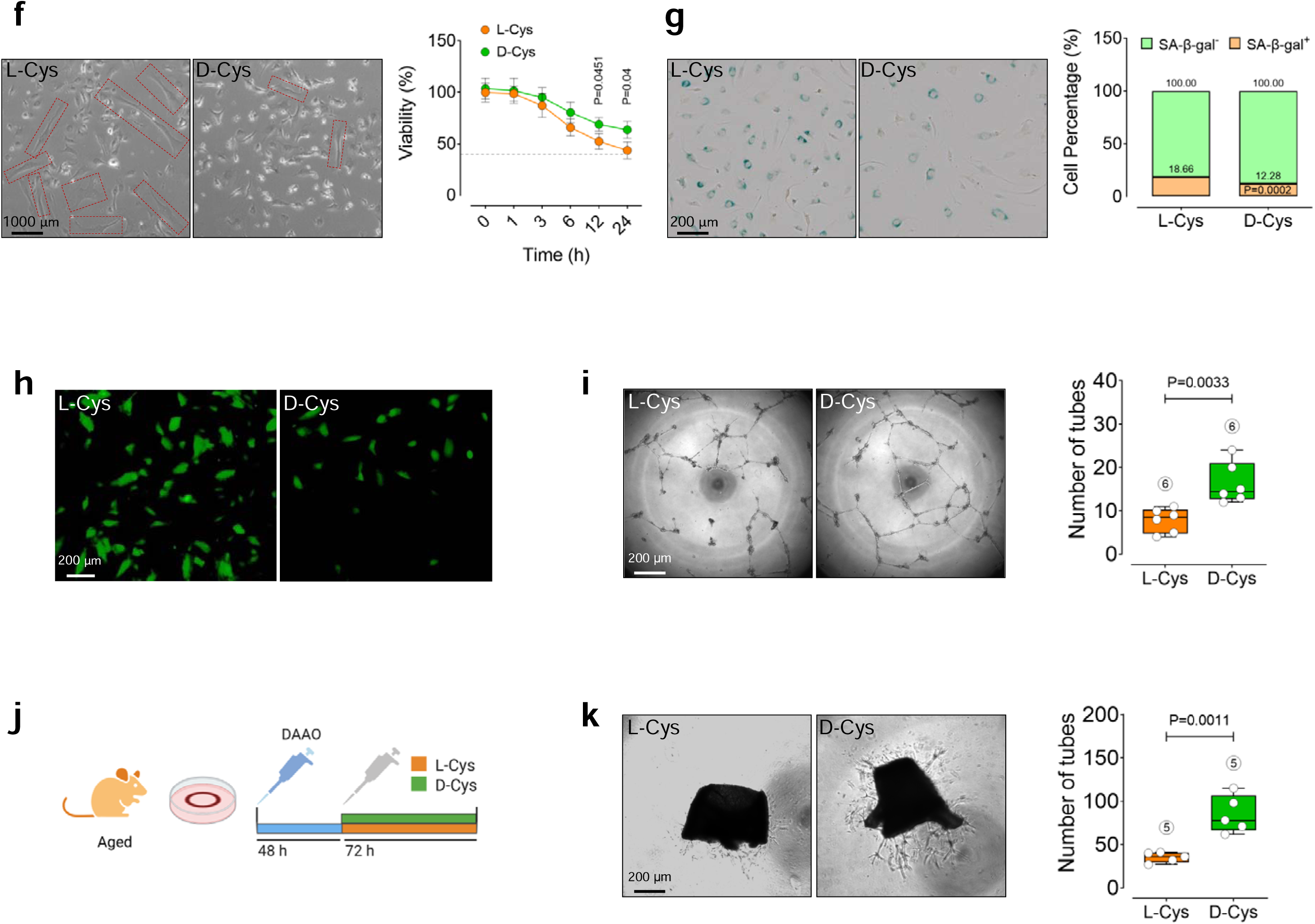
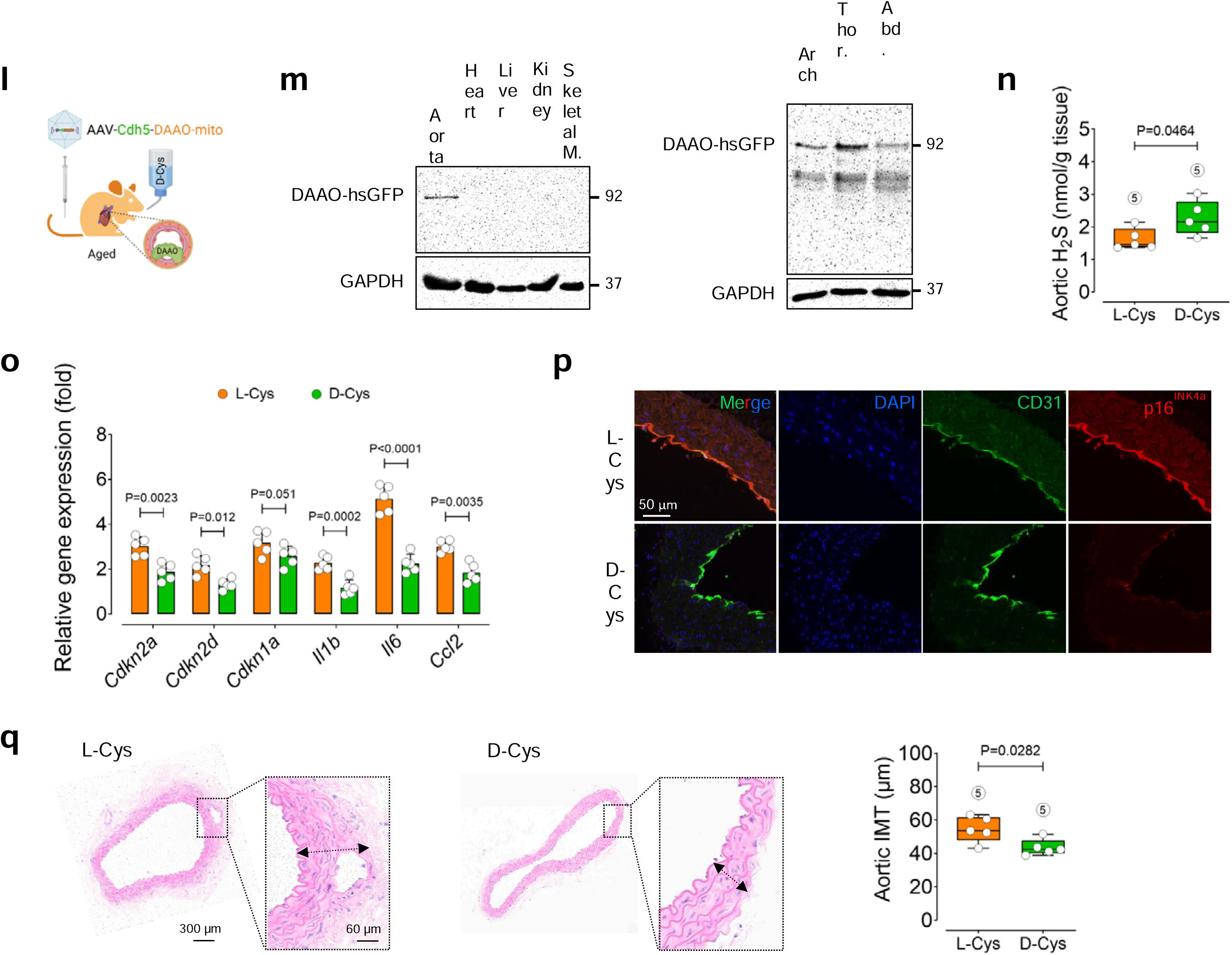
Chemogenetic H_2_S rescues vascular senescence. **a,** Schema for chemogenetic H_2_S production by DAAO. DAAO oxidizes D-cysteine to 3-MP, which is further converted to H_2_S and pyruvate as a byproduct by the endogenous enzyme 3-MPST. H_2_S can be detected by the genetically encoded biosensor hsGFP. **b,** Representative widefield hsGFP images of HAECs co-transfected with mitochondria-targeted DAAO and hsGFP plasmid constructs in the presence of 10 mM L-Cys or D-Cys. **c,** Curves demonstrate significantly higher H_2_S responses in the mitochondria of endothelial cells co-transfected with DAAO and hsGFP constructs following treatment with D-Cys (*n*=22) compared to L-Cys (*n*=20). **d,** Schematic of experimental setting: HAECs (p.15-17) were transduced with Lenti-DAAO-mito for 48 h, followed by treatment with 10 mM L-Cys or D-Cys for 72 h. **e,** Representative immunoblots and quantification of intensities for senescence hallmarks p16^INK4A^, γ-H2A.X, and IL-6. **f,g,** Representative bright-field images depicting senescence phenotype (enlarged, flattened, and multinucleated) (Left) and cell viability percentage (%) (Right) (**f**) and SA-β-gal^+^ cells, confirming senescence-associated lysosomal alterations. Quantitative plot show percentage (%) of SA-β-gal^+^ *vs.* SA-β-gal^−^ cells (**g**) (*n*=6). **h,** CellRox Green staining demonstrates intracellular ROS generation (*n*=6). **i,** Confocal micrographs depict the effects of mitochondrial H_2_S on endothelial angiogenic capacity, represented by numbers of tubes formed (*n*=6). **j,k,** Aortic rings from aged mice were transduced with Lenti-DAAO-mito for 48 h, followed by incubation with 1 M L-Cys or D-Cys for 72 h. Confocal micrographs of aortic endothelial sprouts (*n*=5). **l,** Schematic of experimental setting: Aged mice were infected with an AAV9-Cdh5-DAAO-hsGFP-mito construct and treated with 1 M D-Cys or L-Cys for 4 weeks. **m,** Representative immunoblots for DAAO-hsGFP expression in multiple tissues (Left) and three aortic segments, including arch, thoracic, and abdominal aortas (Right). **n,o,** H_2_S concentrations in aortic tissues (**n**) and transcriptional alterations of CDK inhibitors and SASP components in aortic ECs (**o**) from DAAO-infected aged mice treated with L-Cys or D-Cys (*n*=5). **p,** Immunostaining of aortic ECs for p16^INK4a^ (*n*=5). **q,** H&E staining (left) and intima-media thickness (right) of aortas from these mice (*n*=5). Data were determined in 8-10 micrographs and represent triplicated biologically independent experiments. Scale bars, 20, 30, 50, 60, 100, 200, 1000 μm (**b,f,g,I,k,p,q**). Error bars represent SD (**e,f,g,i,k,n,o,q**) or SEM (**c**). Statistical analysis was performed using two-tailed Mann–Whitney *U*-test (**c**) and two-tailed unpaired Student’s *t*-test (**e,f,g,i,k,n,o,q**). Images created with BioRender.com (**a,d,j,l**).

Given the emerging importance of mitochondrial H_₂_S in regulating cellular senescence^6^, we next tested whether selective restoration of mitochondrial H_₂_S could rescue senescence phenotype. D-Cys administration of Lenti-DAAO-mito-transduced senescent HAECs promotes a robust downregulation of cell-cycle arrest marker p16^INK4a,^ DNA damage response (DDR) marker γ-H2A.X, and the SASP component IL-6 (Fig. 2d,e). Morphologically, treated cells exhibited reversal of characteristic senescence signature (enlarged, flattened, and multinucleated appearance and increased granularity), and restored cell viability (∼63.6% *vs.* ∼43.7%, controls) (Fig. 2f). These results were further confirmed by SA-β-gal activity and ROS generation (Fig. 2g, h). Although D-Cys oxidation by DAAO generates H_2_O_2_ as a byproduct, supplementary experiments demonstrated that exogenous H_₂_S donor NaHS could rescue H_2_O_2_-induced premature senescence (Supplemental Fig. 3), supporting a dominant anti-senescence role of H_2_S in our experimental setup. Our prior studies showed endothelial senescence compromises angiogenic competence^1,2^. We therefore tested whether chemogenetic H_2_S generation restores endothelial angiogenic function. Our data showed the transition of senescent ECs toward an angiogenic phenotype, evidenced by increased numbers of tubes formed (Fig. 2i). As a proof of concept, D-Cys significantly enhanced numbers of endothelial sprouts in aortic rings from aged mice *ex vivo* (Fig. 2j,k). These findings establish that our chemogenetic tool not only profoundly suppresses key senescence hallmarks and acts as a senomorphic intervention but also restores functional properties of senescent ECs.

We next examined whether this strategy could reverse vascular ageing phenotype *in vivo*. Aged mice were infected intravenously with an endothelial-specific adeno-associated viral vector encoding mitochondria-targeted DAAO under the control of the Cadh5 promoter (AAV9-Cdh5-DAAO-mito) (Fig. 2l), led to robust and selective DAAO expression in the aorta with minimal expression in other organs (Fig. 2m). Endothelial-specific induction of mitochondrial H_₂_S generation resulted in significant elevations of H_2_S levels in aortic tissues (Fig. 2n). Of note, no toxic or metabolic changes were seen in treated mice (Supplemental Fig. 4). This intervention produced marked reductions in the expression of senescence hallmarks in aortic ECs, including CDK inhibitors *Cdkn2a* and *Cdkn2d*, as well as the SASP components *Il1b*, *Il6*, and *Ccl2* (Fig. 2o,p). Importantly, this intervention also mitigated aortic tissue remodeling in aged mice, evidenced by reduced intima-media thickness (IMT) (Fig. 2q) and downregulated ECM remodeling markers *Col1a1, Mmp2*, and *Mmp9* (Supplemental Fig. 5), indicating H_₂_S-dependent improvement of aged vascular wall structure.

Collectively, these findings identify age-associated loss of endogenous H_₂_S production as a conserved feature of vascular senescence and atherosclerosis. More importantly, we demonstrate that this process is reversible. Our findings provide direct evidence that impaired H_₂_S signaling is not merely a passive biomarker of vascular ageing but represents a modifiable driver of vascular senescence and remodeling. This work introduces a remote-control chemogenetic H_₂_S donor system that enables precise cell-specific, mitochondria-targeted H_₂_S generation with added advantage of temporal on–off switching. Unlike traditional H_₂_S donors, this strategy allows targeted restoration of H_₂_S signaling in endothelial cells with least off-target effects, offering a powerful experimental and potentially translational platform for vascular ageing. In light of recent evidence linking H_₂_S deficiency to cardiometabolic disease^17,23^ and reduced healthspan^9^, these findings position our chemogenetic system as a potential senotherapeutic tool for combating vascular ageing and its cardiovascular consequences.

## Methods

### Human cohorts

#### ASCVD cohort

We obtained plasma samples from aged patients with ASCVD (*n*=110; >80 years old) enrolled in *Aging Heart Zurich Cohort—* an ongoing observational prospective human cohort conducted at University Hospital Zurich^24^ (Supplementary Table 1). All patients were diagnosed for ASCVD, including atherosclerosis, coronary artery disease, and arrhythmias, at baseline in 2020 and are followed up every 1 year. Atherosclerosis was assessed by imaging studies cardiac CT scan and coronary angiography. Fasting blood test was performed for biochemical parameters including blood glucose, total cholesterol, LDL-cholesterol, HDL-cholesterol, triglyceride, Hemoglobin A1c (HbA1c), CRP, BUN, creatinine, alanine aminotransferase (ALT), aspartate aminotransferase (AST), and gamma-glutamyl transferase (GGT), troponin T, and NT-proBNP. All participants received no compensation and provided written informed consent. The study was approved by the Kantonale Ethikkomission Zurich (BASEC-Nr: 2020-02597). Plasma samples from a separate cohort of 77 healthy controls were obtained from the blood bank of the Swiss Red Cross.

#### Aging human study

To validate the associations between plasma H_2_S levels and age in healthy humans, we obtained plasma samples from a cohort of 86 healthy young (*n*=41; 18-40 years old) and aged (*n*=45; >65 years old) subjects from the blood bank of the Swiss Red Cross and processed accordingly.

#### Validation human study

CAD Patients (*n*=5) were undergone to coronary artery bypass graft (CABG) surgery at Department of Cardiac Surgery, University Hospital Zurich, Switzerland (Supplementary Table 2). All procedures were approved by the local ethics committee and performed in accordance with the Declaration of Helsinki (Kantonale Ethikkommission Zürich, BASEC number: 2023-02040). Plasma samples were obtained and stored at -80 C for further H_2_S quantification. Samples from non-CAD transplant donors (*n*=5) served as controls.

### Mouse models

Wild-type 8-week old C57BL/6J female and male mice were purchased from the Jackson Laboratory and aged in the animal facility of University of Zurich, Schlieren. Accordingly, mice were fed a standard chow diet (containing 19% Protein, 61% Carbohydrate and 7% Fat, #D11112201, Research Diets, New Brunswick, NJ, USA) until 3 (as Young) or >24 (as Old) months of age. Our study complies with all ethical regulations approved by the Institutional Animal Care and Use Committee at University of Zurich and the Cantonal Ethics Commission Zurich (ZH023/2023 and ZH241/19).

Atherosclerosis-prone *Ldlr*^−/-^ C57BL/6J mice were fed a specific Western diet (21% fat, 0.21% cholesterol, SSNIFF TD. 88137, E15775-34; Supplementary Table 3) for 12 weeks, as indicated in each experiment. Mice were kept at the animal facility of the University of Milan, Italy. All animal procedures were approved by the pertinent Ethical Committees (Autorizzazione Ministeriale 384/2019-PR, 92/2020, 240/2021 and 565/2022).

Animal care and all experimental protocols were in accordance to the principles of the ‘3Rs’—replacement, reduction and refinement—according the Directive 2010/63/EU of the European Parliament and of the Council of 22 September 2010 on the protection of animals used for scientific purposes. All mice were housed in controlled environment in plexiglass cages under a strict 12 h:12 h light/dark cycles at an ambient temperature (23 ± 1°C) and humidity of 55 ± 10%, and had free access to drinking water and food *ad libitum*.

### Construction and chemogenetic DAAO activation *in vivo*

The DAAO-mito construct (a mitochondria-targeted version of DAAO, as described previously^19^) was packaged into an adeno-associated virus (AAV) expression vector (Stratagene) with a vascular endothelial-cadherin 5 (Cdh5) promoter. The Cdh5 has been extensively characterized as an endothelial cell-specific promoter. The viral construct was packaged into AAV9 viral particles via Rep2/Cap9 AAV encapsulation construct, as previously described^25,26^. ∼1×10^12^ viral genome copies were suspended in 100 µL of PBS and injected via the tail vein into aged mice. To activate DAAO, D-Cysteine (Santa Cruz Biotechnology, sc-255054D) was added to the drinking water at a final concentration of 1 M for 4 weeks.

### Blood and tissue harvesting

Blood samples were obtained by cardiac puncture to further collect plasma by centrifugation at 125 x g (8 min) at room temperature and frozen at −80 °C^1^. After blood collection, heart was perfused for aorta harvesting.

### H_2_S level quantification

H_2_S levels in plasma and aortic tissues from human and mouse subjects were measured by gas chromatography—chemilumunescence (Agilent Technologies, Santa Clara, CA, USA), as previously described^17,23^.

### Blood lipid profiling

Total cholesterol and triglycerides were quantified in plasma from wild-type and atherosclerosis-prone mice by colorimetric enzymatic technique (ABX Pentra, HORIBA Medical) according to the manufacturer’s protocols.

### Serum creatinine and BUN measurement

Serum creatinine and BUN were measured using colorimetric detection kits (Cayman Chemical, 700460 and Invitrogen, EIABUN, respectively), according to manufacturers’ instructions. Absorbance was read at 495 nm and 450 nm for creatinine and BUN, respectively, with a Tecan microplate reader.

### Plasma ALT and AST measurement

Plasma alanine aminotransferase (ALT) and aspartate aminotransferase (AST) were measured using colorimetric detection kits (Cayman Chemical, 700260 and 701640, respectively), according to manufacturers’ instructions. Absorbance was read at 340 nm for both ALT and AST with a Tecan microplate reader.

### Cell lines

HAECs (Lonza, CC-2535) were cultured in EBM-2 medium (Lonza, 00190860) supplemented with EGM-2 (10% FBS, 2 mM L-glutamine, 100 μg/mL penicillin-streptomycin, Lonza, CC-3162/6) at 37°C, 5% CO_2_. Cells at passages 4–6 were used as proliferating controls, whereas cells at passages 15–17 represented the replicative senescence state. Cellular senescence was confirmed by proliferative arrest (reduced cell count), SA-β-gal staining, and p21^WAF1/Cip1^ immunoblots, as previously described^1,2^.

### Cellular senescence models

#### Replicative senescence

Proliferating HAECs (p.4-6) were cultured in EGM-2 (5,000 cells/cm²) and passaged every 48 h until the proliferation is arrested and senescence phenotype is confirmed. As previously established, cellular senescence was verified by SA-β-gal staining and marked reduction in cell count (stained with 0.4% Trypan Blue)^1,2^.

#### H_2_O_2_-induced premature senescence

Proliferating HAECs at passages 4-6 were cultured in EGM-2 (+10% FBS), respectively, followed by treatment with exogenous H_2_O_2_ (50 μM) for 4 h. The culture media were then replaced, and cells were maintained for up to 72 h.

### Lentiviral DAAO transduction

The mitochondria-targeted version of DAAO containing the corresponding signal sequence was cloned and packaged using a mammalian lentiviral system (Addgene plasmids #12259, #12260), as described previously^1,2^. HAECs (p.15-17) were grown to ∼70-80% confluence and transduced with lentivirus-DAAO-mito (multiplicity of infection 20–50) in the serum-free medium, replaced with L-Cysteine-free, 10% FBS medium after 5 h. 48–72 h post-transduction, cells were treated with 10 mM L-Cysteine (Santa Cruz Biotechnology, sc-286072B), as control, or D-Cysteine for 48 h to generate mitochondrial H_2_S and subsequently subjected to senescence phenotype analysis.

### hsGFP fluorescence imaging

∼70-80% confluent HAECs were co-transfected with the DAAO-mito plasmid (pC1-CMV-mito-DAAO, a gift from Vsevolod Belousov, Addgene plasmid #141132; http://n2t.net/addgene:141132; RRID:Addgene_141132) and the H_2_S biosensor hsGFP plasmid (pcDNA3-hsGFP, a gift from Huiwang Ai, Addgene plasmid #64912; http://n2t.net/addgene:64912; RRID:Addgene_64912) using PolyJet transfection reagent (SignaGen Laboratories, SL100688)^12,13^. After 5 h, the serum-free culture medium was replaced with L-cysteine-free, 10% FBS medium. 24 h post-transfection, coverslips were mounted on a live-cell imaging platform and incubated with 10 mM L- or D-Cys in HEPES-buffered solution (pH 7.4) at 37 °C for real-time fluorescence imaging. The hsGFP biosensor was excited at 480 nm, with emission captured at 525 nm using Metafluor Software (Molecular Devices)^21,22^. Mitochondrial H_2_S was quantified as the mean fluorescence intensity (F_Intensity_).

### Co-transfection of stDCyD—hsGFP

HAECs were grown to ∼70-80% confluence and co-transfected with the DsRed.stDCyD and the hsGFP (pcDNA3-hsGFP, a gift from Huiwang Ai, Addgene plasmid #64912; http://n2t.net/addgene:64912; RRID:Addgene_64912) plasmids^12^ using PolyJet transfection reagent in the serum-free culture medium for 5 h. Culture media were replaced by L-cysteine-free, 10% FBS medium. 24 h post-transfection, coverslips were mounted on a live-cell imaging platform and incubated with 10 mM L- or D-Cys in HEPES-buffered solution (pH 7.4) at 37 °C for real-time fluorescence imaging.

### Widefield epifluorescence microscopy

Widefield epifluorescence microscopy was conducted using the Axio Observer Z1.7 inverted system (Carl Zeiss AG, Oberkochen, Germany), featuring a high-resolution Plan-Apochromat 40×/1.4 numerical aperture oil immersion lens and a monochrome Axiocam 503 CCD camera. A custom-built perfusion apparatus was integrated to enable precise delivery and removal of experimental solutions. Cells expressing DsRed-stDyCD were illuminated with 555/30 nm light, and emitted fluorescence was filtered through an FT570 dichroic mirror and 605/70 nm bandpass filter. Detection of H_2_S levels with the hsGFP biosensor was achieved by exciting at 480 nm and capturing emission at 525 nm. Image acquisition and system control were executed using Zen Blue 3.1 Pro software (Carl Zeiss AG).

### Senescence Associated-β-galactosidase (SA-β-gal) assay

SA-β-galactosidase staining was performed using Cellular Senescence assay kit (Merck Millipore, KAA002), as previously described^1,2^. Briefly, cells were fixed and stained with x-gal solution overnight at 37°C. Senescent cells, illuminated as blue-stained cells, were captured under an Agilent BioTek Lionheart FX automated confocal microscope and analyzed using ImageJ software 1.54j (Fiji).

### Cell viability test

Cultured cells were trypsinized, resuspended in culture medium, and mixed 1:1 with 0.4% Trypan Blue solution, as previously described^1,2^. After 3 min incubation at room temperature, viable (unstained) and non-viable (blue-stained) cells were counted under a light microscope. The percentage of viable cells was finally reported.

### CellRox Green staining

Cells were grown to ∼70-80% confluence and treated with CellROX Green reagent at a final concentration of 5 μM (Invitrogen, C10444) for 30 min at 37 °C to detect intracellular ROS by an Agilent BioTek Lionheart FX automated confocal microscope. The fluorescence intensity was evaluated using ImageJ 1.54j.

### Real-time Quantitative PCR

Total RNA from cells and tissues was extracted using TRIzol Reagent^®^ (Sigma-Aldrich, T9424), as previously described^1,2,26,27^. Quantitative PCR was performed using the Power SYBR Green PCR Master Mix (Thermo Fisher Scientific, 4367659) on a Quant Studio 5 and 7 cyclers system (Life Technologies, ThermoFischer Scientific, Switzerland) according to the manufacturer’s instructions. The sequences of all primers used for qPCR are listed in Supplementary Table 4.

### Immunoblotting

Cells and tissues were lysed in ice-cold RIPA buffer (ThermoFisher Scientific, 89901) or nuclear and cytoplasmic extraction reagents (ThermoFisher Scientific, 78835) supplemented with Halt protease and phosphatase inhibitors cocktail (ThermoFisher Scientific, 78442). Total protein was quantitated using the BCA protein assay kit (ThermoFisher Scientific, 23225). Equal amounts of proteins (15-20 µg) were separated on pre-made gels (8-12%, BoltTM Bis-Tris Plus mini protein gels 1.0 mm, ThermoFischer Sceintific, NW00100BOX) in a SDS-PAGE gel electrophoresis system (mini blot module, ThermoFischer Scientific, B1000), followed by electroteranfer and further incubation with specific antibodies. The protein bands were visualized using Amersham Imager 600 (GE Healthcare) and quantitative densitometric analyses were performed using ImageJ software 1.54j (Fiji).

All antibodies used in immunoblotting are listed in Supplementary Table 5.

### Immunostaining

Aortas and ECs were fixed in Bouin’s fixative (Muto Pure Chemicals) or 4% paraformaldehyde (Santa Cruz Biotechnology, sc-281692) for 2 h at room temperature^28^, followed by paraffin-embedding. Tissues were then sectioned (5-μm), deparaffinized, and blocked with Streptavidin/Biotin blocking solution (Vector Laboratories) and 5% goat serum. Slides were stained with CD31, p16^INK4a,^ or Ki67 primary antibodies (1:100) for 1 h at room temperature, followed by washing with 1x PBST for 3 times. Slides were further incubated with the respective secondary Fluor chrome antibodies (conjugated with AlexaFluor 488 or 594; 1:200) for 30 min at room temperature, and further incubated with mounting media containing DAPI (for nuclear counterstaining) for scanning under fluorescent confocal microscope (Leica TCS-SP8).

### Aortic root histology

Aortas were collected from wild-type and *Ldlr^−/−^* mice fed a chow diet or Western diet. For aortic root histological analysis, tissues were fixed in 4% paraformaldehyde overnight, paraffin-embedded, and sectioned at 5 µm thickness. Sequential sections were stained with H&E for plaque area analysis. The sections were dehydrated through graded ethanol and cleared in xylene. Images were acquired using a Zeiss confocal microscope using the Zen software and analyzed with ImageJ software 1.54j (Fiji). Plaque area was measured on H&E-stained sections.

### Aortic H&E staining

Aortas were collected from DAAO-infected aged mice for H&E staining and histological analysis. Tissues were fixed in 4% paraformaldehyde (Santa Cruz Biotechnology, sc-281692) overnight, paraffin-embedded, and sectioned at 5 µm thickness. Sequential sections were stained with H&E for IMT analysis. The sections were dehydrated through graded ethanol and cleared in xylene. Images were acquired using a Zeiss confocal microscope using the Zen software and analyzed with ImageJ software 1.54j (Fiji). Aortic IMT was analyzed by QuPath v0.6.0 software.

### Endothelial tube formation assay

Tube formation assay was performed to analyze angiogenic capacity of ECs, as previously described^1,2^. Cells were cultured onto Matrigel (Corning 356234) at a subconfluent level. Real-time images were recorded every 15 min for 16 h at ×6.4 magnification using a phase-contrast Olympus IX81 microscope (Hamamatsu C11440 detector, 1 megapixel, 16 bit). Tube formation was analyzed using ImageJ Angiogenesis Analyzer (v.1.54j).

### *Ex vivo* aortic ring sprouting assay

Aortic ring sprouting Aortas from aged mice, cut into 2-mm fragments, and cultured onto Matrigel. Aortic rings were then transduced with Lenti-DAAO-mito (multiplicity of infection 20–50) in the serum-free medium, replaced with L-Cysteine-free, 10% FBS medium after 5 h. 48 h post-transduction, aortic rings were treated with 1 M L- or D-Cysteine for 72 h. We acquired real-time images daily (for 5 days) using an incubator-equipped phase-contrast Olympus IX81 microscope with the Hamamatsu (C11440) detector, followed by the analysis of the number of aortic sprouts^62^.

### *Ex vivo* vasorelaxation assay

Murine aortic fragments (2-mm) were mounted in a 5-ml organ chamber filled with Krebs-Ringer bicarbonate solution (37 °C, pH 7.4, bubbled with 95% O_2_/5% CO_2_) and connected to an isometric force transducer (PowerLab 8/30 and LabChart v.7.2.5, AD Instruments) for continuous isometric tension recording (Multi-Myograph 610M, Danish Myo Technology). Concentration-response curves were obtained in responses to increasing concentrations of acetylcholine (Ach, 10^−9^ to 10^−5^ M; Sigma-Aldrich, A6625)^66^.

### Statistics and reproducibility

No statistical methods were used to pre-determine sample sizes, but similar sample sizes were used in previous publications^1,2,28–30^. In the ASCVD cohort and the validation human study, no sample size calculation was conducted, as samples were retrospectively collected from observational studies. The cohorts were assembled with the goal of maximizing biological and phenotypic depth rather than targeting a predefined effect size. Continuous variables were checked for normality and compared using unpaired two-tailed Student’s *t*-test. No method of randomization was used to assign animals to experimental groups. No animals or data points were excluded from experiments or analyses. Animals were monitored daily for sickness symptoms and body weight during their lifespan. They were euthanized immediately at the clinical end point when recommended by veterinary and biological service staff members. The investigators in this study were blinded to the conditions of the experiments. *n* represents the number of samples or animals used. At least three independent triplicated experiments were performed for each experimental set-up. Data are expressed as mean ± SD or SEM, and *P*<0.05 was defined as statistically significant. Continuous data were shown as median ± SD or interquartile ranges (IQR) if skewed. Statistical analysis was performed using GraphPad Prism 9 (v.9.5.1, La Jolla, CA, USA). Comparisons between groups were conducted using unpaired two-tailed Student’s *t*-test and one-way or two-way ANOVA with Tukey’s post *hoc* test. Spearman’s rank-order correlations were used to analyze associations between plasma H_2_S levels and either aortic root lesion area (%) or vasorelaxation (%), respectively.

## Data availability

Source Data and Supplementary Information are provided with this paper. The human study data used in this research are maintained by the Departments of Cardiology and Cardiac Surgery at University Hospital Zurich and are not publicly available due to patient privacy and data protection regulations. Researchers seeking access must apply through the established procedures outlined by the Aging Heart Zurich Cohort (https://www.usz.ch/studie/aging-heart-zuerich-beobachtungsstudie-zur-lebensqualitaet-von-ueber-80jaehrigen-patienten-mit-einer-herzerkrankung/). Access is granted upon approval and may require specific ethical and governance clearances. The GC data and processed datasets have been securely stored on an institutional research repository at University of Zurich. These data are available upon request from the corresponding authors and subject to institutional data-sharing policies. All other data and reagents that support the findings of this study are available upon reasonable request from the corresponding author.

## Author contributions

S.S.S.S. conceptualized and designed the study, and drafted, wrote, and revised the manuscript. K.S., T.S., L.D.D., and M.C. performed the experiments, including mouse studies, *in vitro* cellular studies, senescence and signal transduction studies, and real-time fluorescence imaging. K.S. and T.S. performed plasma and tissue H_2_S measurement and analyzed data. M.C., A.A., L.D.D., and G.D.N. conducted premature aging and atherosclerosis-prone *Ldlr*^−/-^ mouse modeling and provided tissues and data. H.R. and O.D. collected samples from non-CAD and CAD patients undergoing CABG surgery. E.E. and A.Z. designed a H_2_S biosensor and conducted complementary real-time fluorescence imaging. S.S.S.S. and M.H. contributed to management and sample collection within Aging Heart Zurich Cohort. S.S.S.S., F.P., and F.R. contributed to interpretation of data. All authors provided intellectual contributions and approved the final manuscript. S.S.S.S. directed and supervised all aspects of the study.

## Acknowledgements

The authors acknowledge funding from the Swiss National Science Foundation Spark grant (#CRSK-3_229134), the Iten Kohaut Foundation in collaboration with the USZ Foundation (#2512), the Peter Bockhoff Foundation (SCWM-280168), Novartis Foundation for Medical-Biological Research (#21A053), and the SwissLife Jubiläumsstiftung (#1565) (to S.S.S.S.). K.S. has received support from SwissLife Jubiläumsstiftung (#1438 and #1563), Wael Almahmeed International Atherosclerosis Society (IAS) Research Fellowship, and GRC Travel Grant from University of Zurich (2025_Q3_TG_075). T.S. is funded by the SwissLife Jubiläumsstiftung (#1564), Wael Almahmeed International Atherosclerosis Society (IAS) Research Fellowship, and GRC Travel Grant from University of Zurich (2026_Q1_TG_173). The Aging Heart Zurich Cohort receives support from VELUX stiftung and Accentus stiftung.

