## Supplemental Figures for "Chemogenetic Hydrogen Sulfide Rescues Vascular Senescence in Ageing"

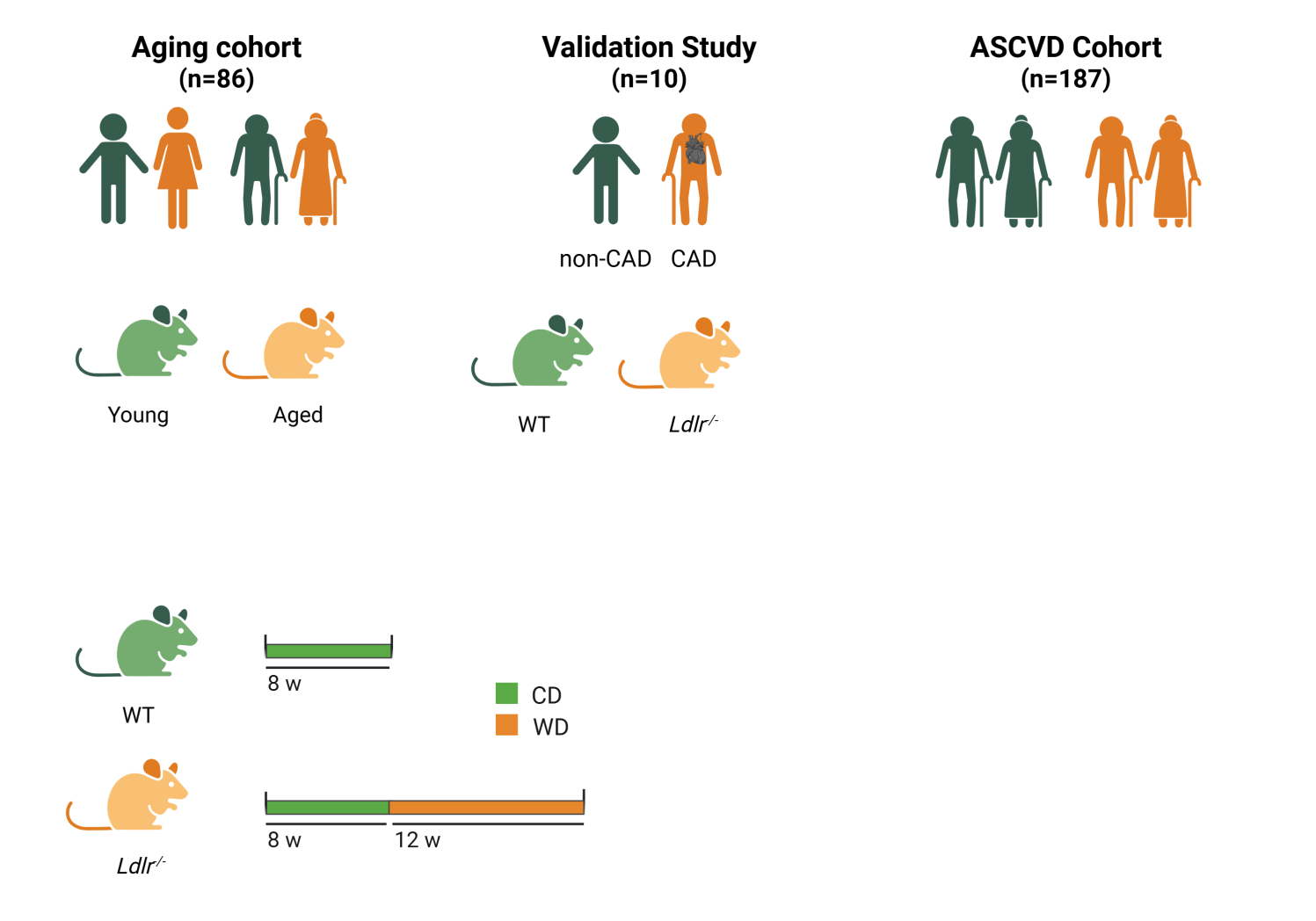
**Supplemental Figures and Legends**

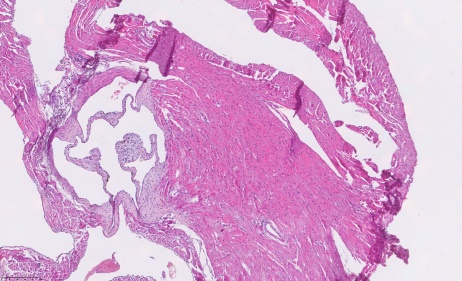

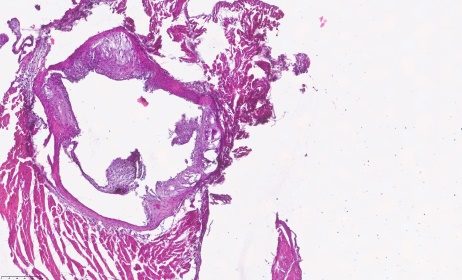

200 μm

WT

*Ldlr^-/-^*

**c**

**d**

**b**

**a**

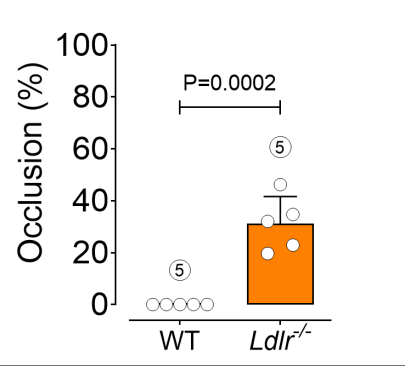

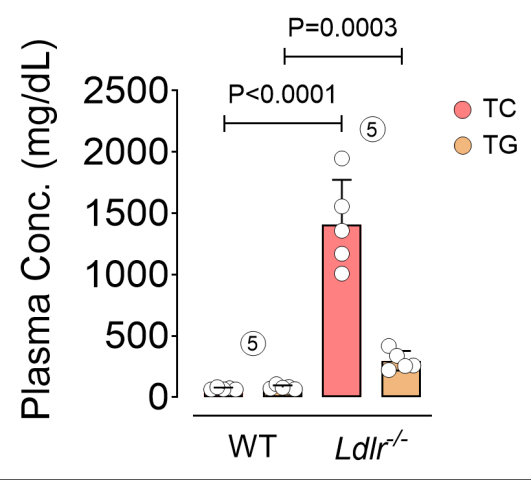

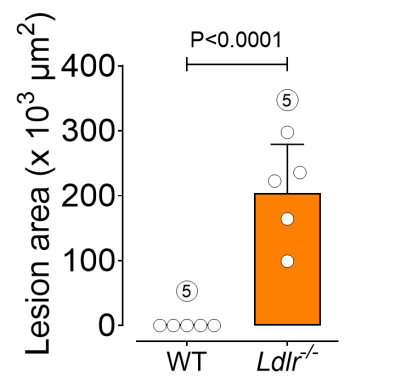

**Supplemental Fig. 1.** **Characterization of atherosclerosis in *Ldlr^-/^*^-^ mice. a,** WT and *Ldlr^−/−^* mice were fed chow (for 8 weeks) or Western diet (for 12 weeks), respectively. **b,c,** Representative images (**b**) and quantification (**c**) of H&E staining of aortic root lesion (x 10^3^ μm^2^) and luminal occlusion (%) (*n*=5). Arrowheads indicate plaque areas. **d,** Total cholesterol (TC) and triglyceride (TG) concentrations in plasma (*n*=5). Data were determined in 8-10 micrographs. Scale bar, 200 μm (**b**). Error bars represent SD (**c,d**). *P* values were calculated using two-tailed unpaired Student’s *t*-test (**c,d**). Images created with BioRender.com (**a**).

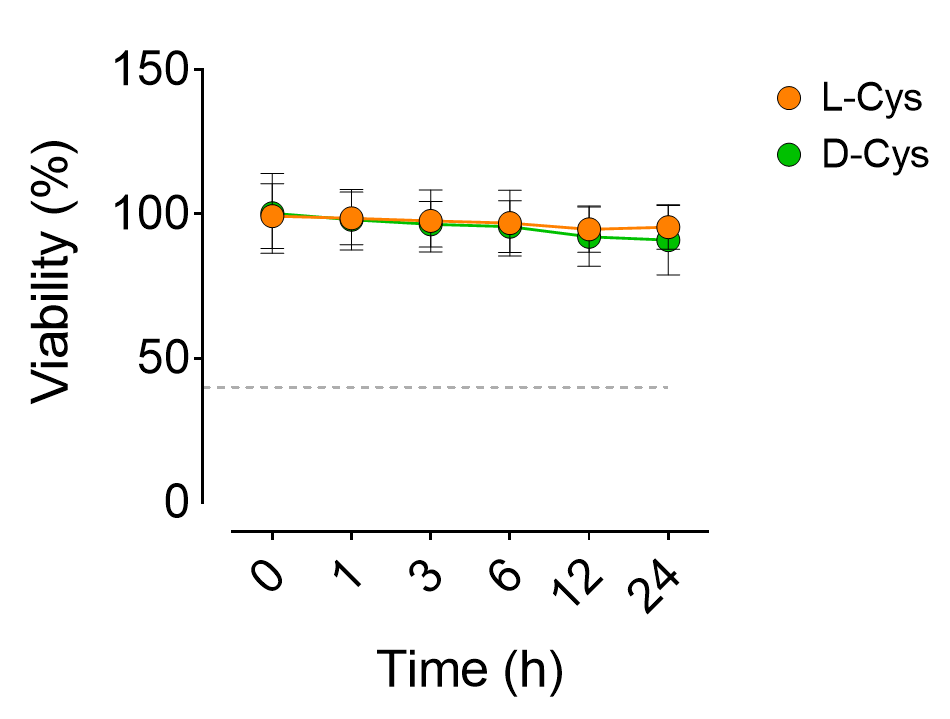

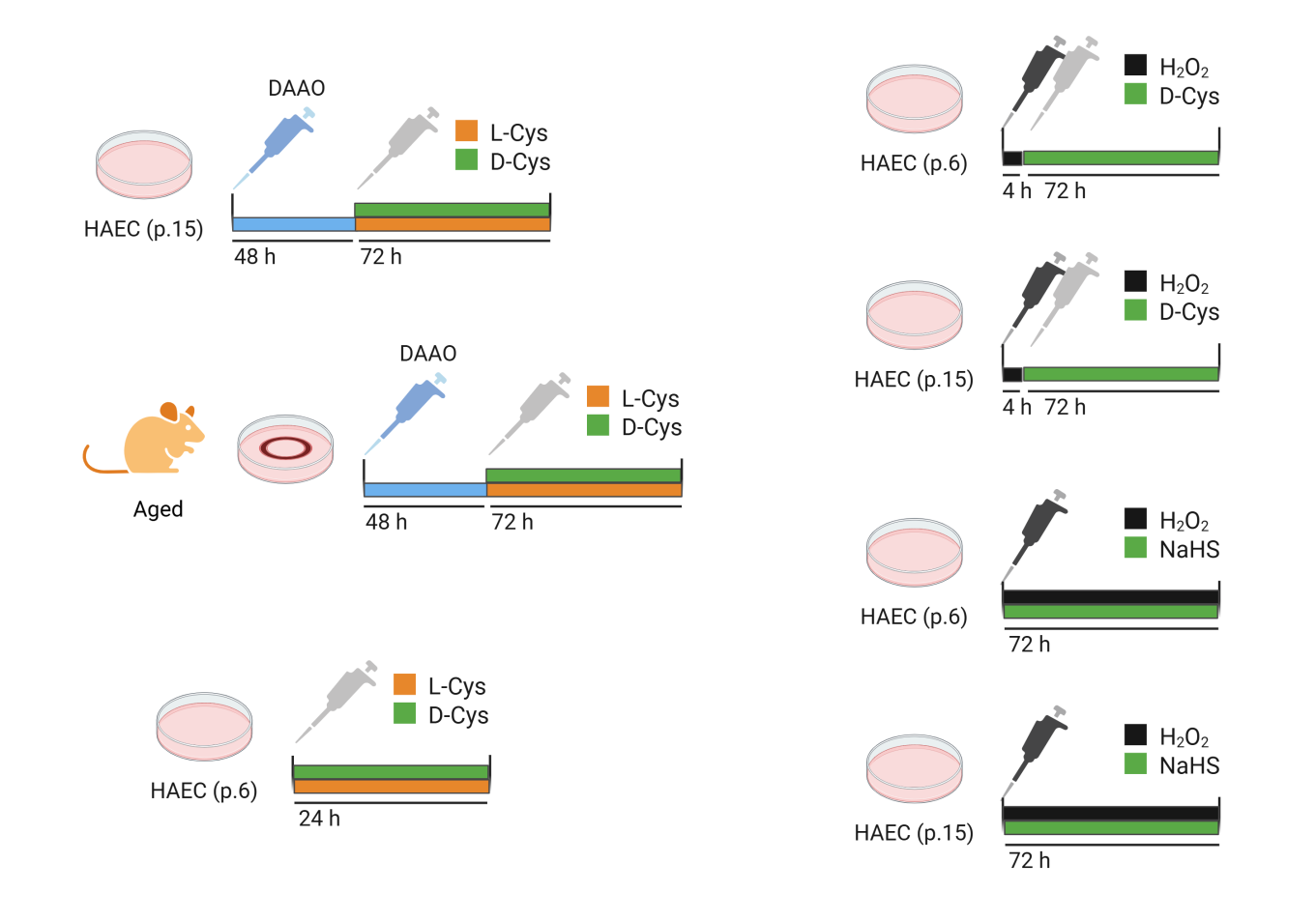

**Supplemental Fig. 2.** **D-Cys is non-toxic in endothelial cells.** HAEC (p.6) were treated with 10 mM L-Cys or D-Cys, followed by cell counting for viability assessment. Curve demonstrates cell viability percentage (%) (*n*=6). Data represent triplicated biologically independent experiments. Error bars represent SD. Statistical analysis was performed using a two-way ANOVA followed by Tukey’s post *hoc* test.

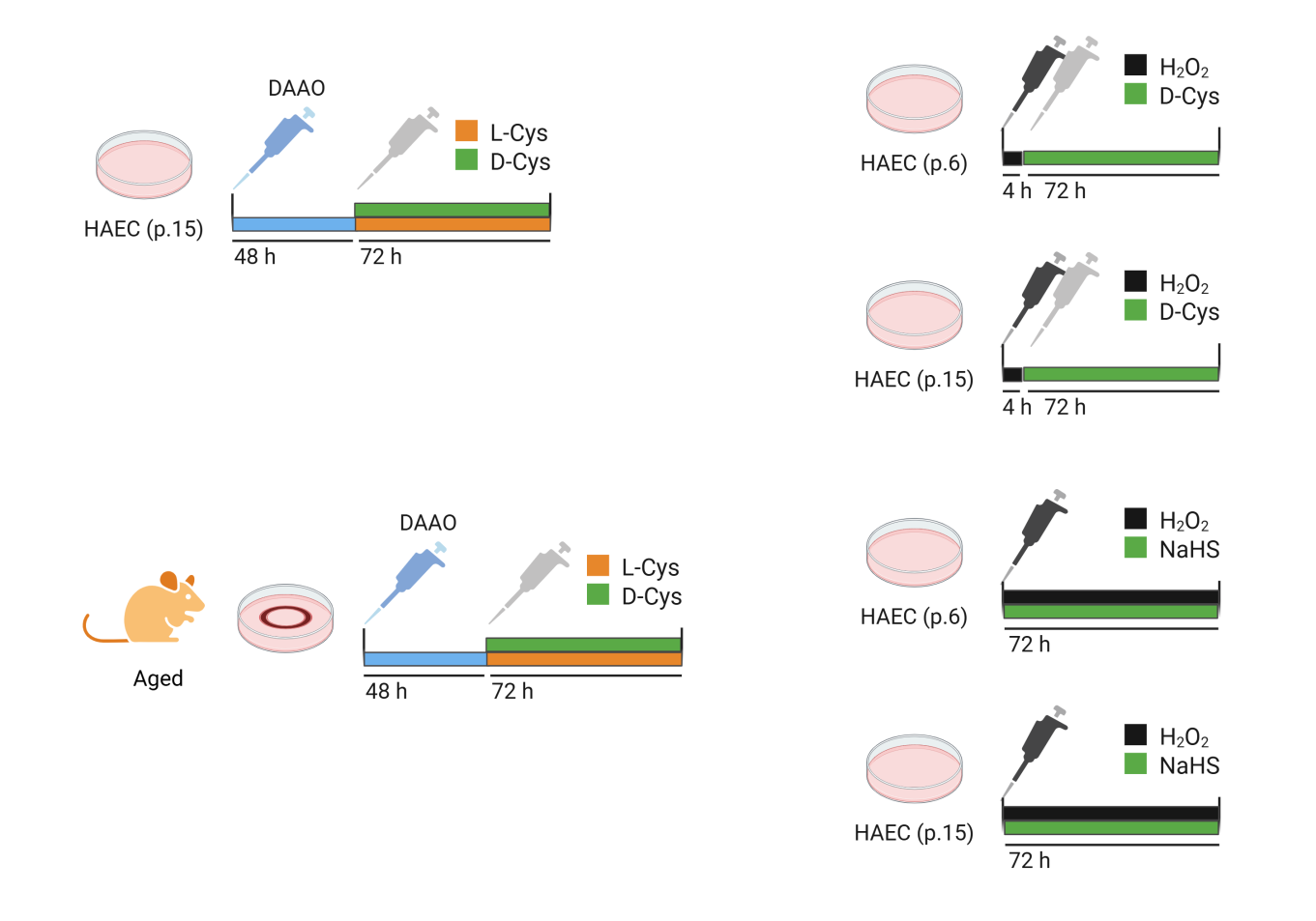

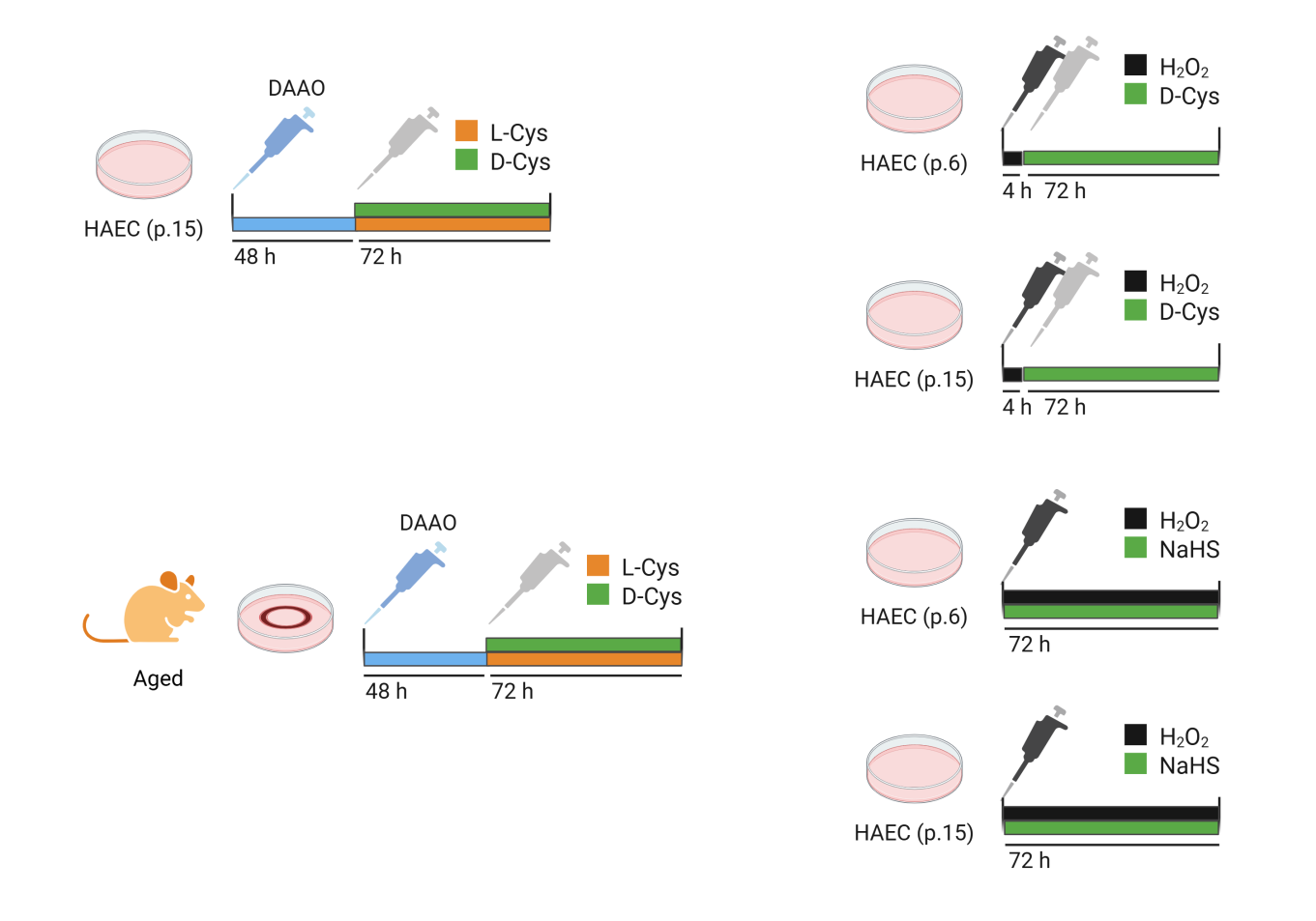

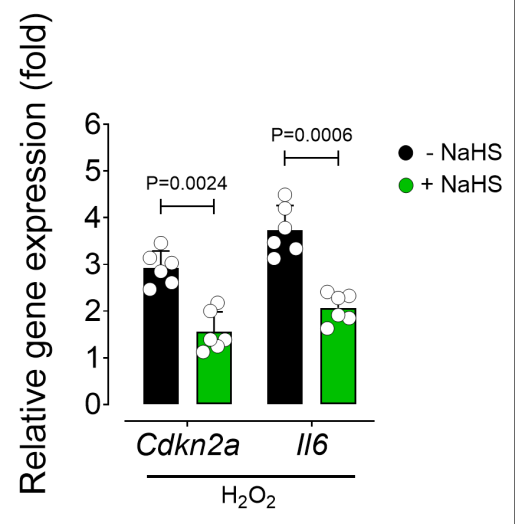

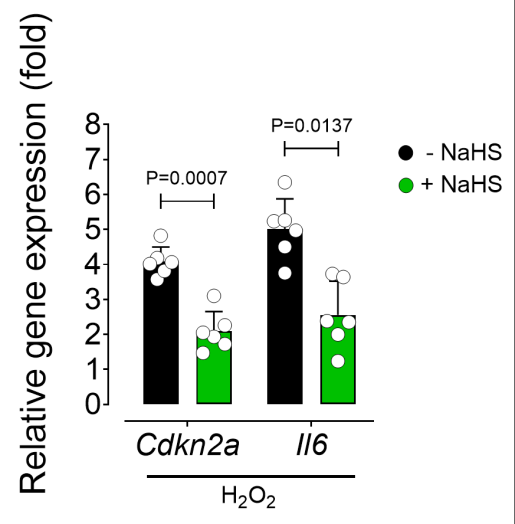

p.15

p.6

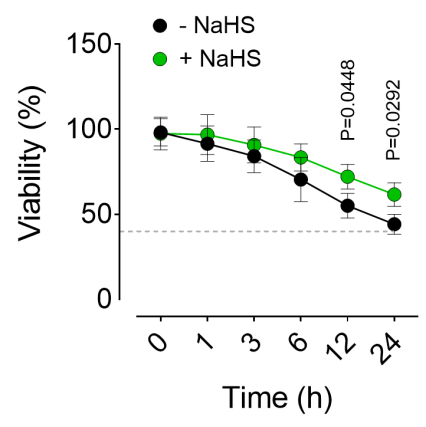

p.15

p.6

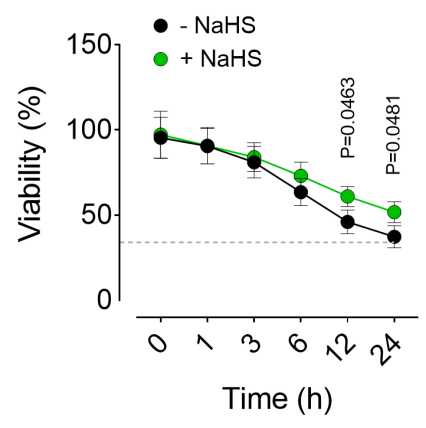

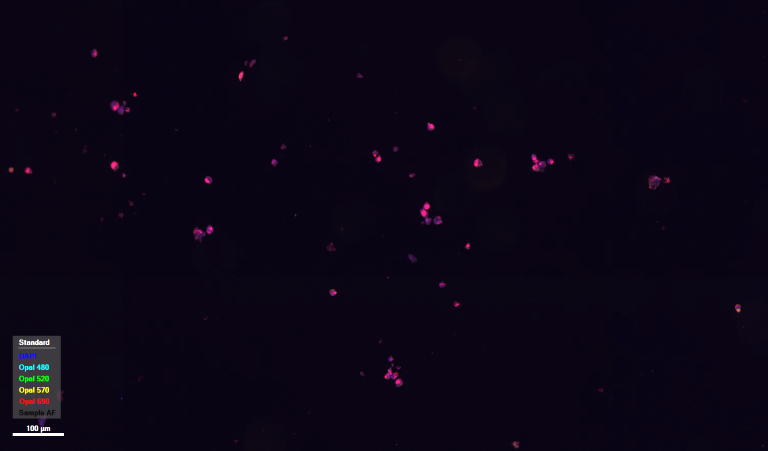

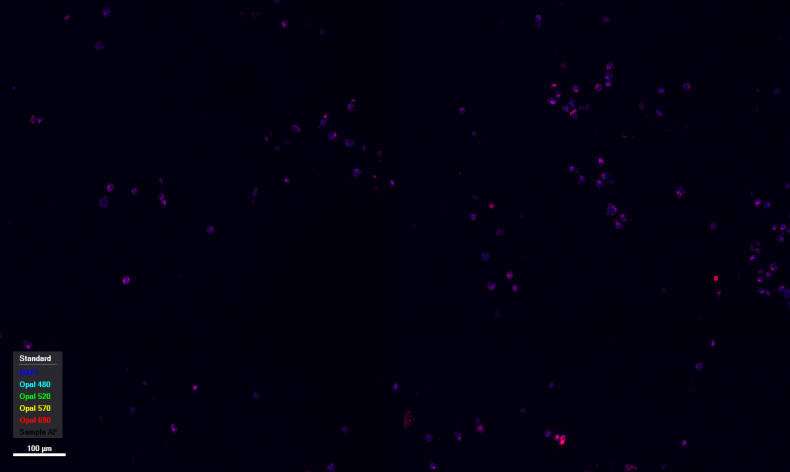

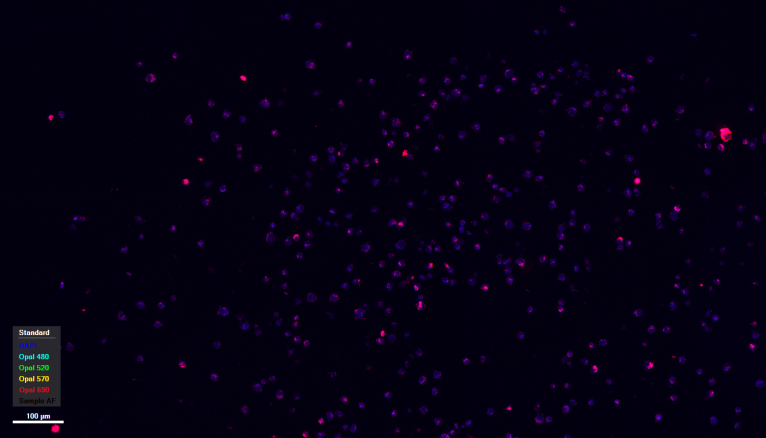

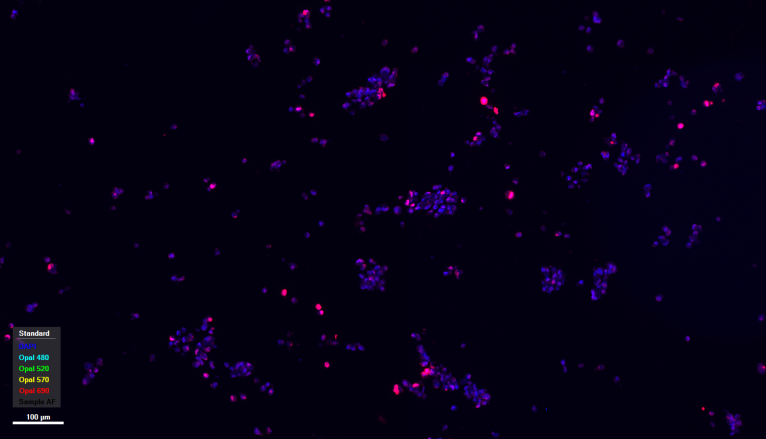

DAPI

Ki67

100 μm

+ NaHS

- NaHS

p.15

p.6

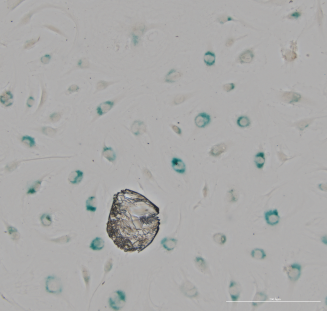

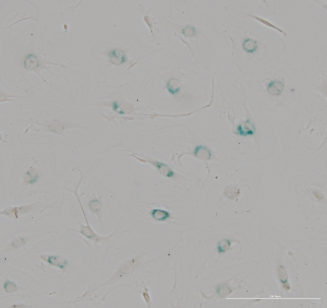

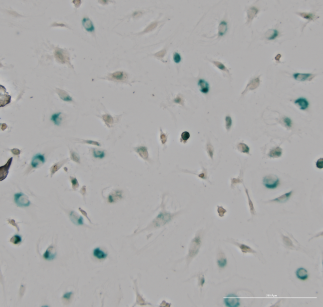

200 μm

+ NaHS

- NaHS

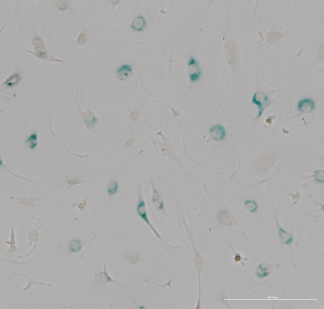

p.15

p.6

**e**

**d**

**c**

**b**

**a**

**Supplemental Fig. 3.** **NaHS rescues H_2_O_2_-induced premature senescence.** **a,** HAEC at lower (p.6) and higher (p.15) passages were co-incubated with NaHS (100 µM) and exogenous H_2_O_2_ (10 µM) for 72 h, followed by senescence phenotype characterization. **b,** qPCR shows transcriptional alterations of the CDK inhibitor *Cdkn2a* and the SASP component *Il6* in these cells (*n*=6). **c,** Representative bright-field images of SA-β-gal^+^ cells. **d,** Immunostaining for the proliferation marker Ki67. **e,** Curve demonstrating cell viability percentage (%) (*n*=6). Data were determined in 8-10 micrographs and represent triplicated biologically independent experiments. Scale bars, 100 and 200 μm (**c,d**). Error bars represent SD (**a,b,e**). Statistical analysis was performed using two-tailed unpaired Student’s *t*-test (**a,b**) and a two-way ANOVA followed by Tukey’s post *hoc* test (**e**). Images created with BioRender.com (**a**).

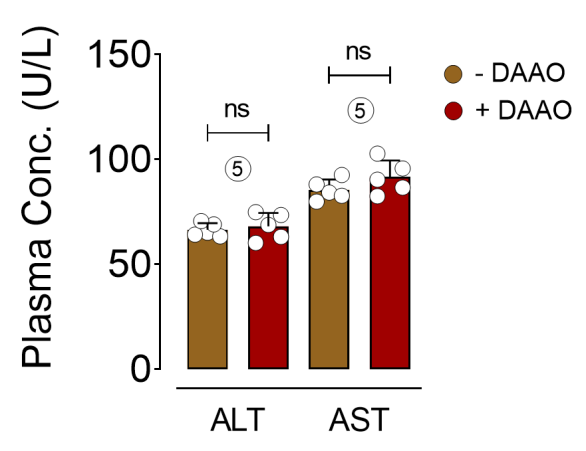

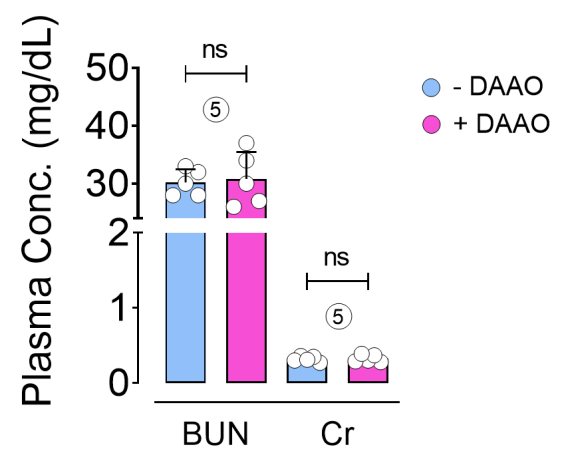

**b**

**a**

**Supplemental Fig. 4.** **Kidney and liver function profiles in DAAO-infected mice. a,** Kidney function profile, represented as serum BUN (Left) and creatinine (Cr) (Right) in aged mice infected with AAV-Cdh5-DAAO construct or vehicle (*n*=5). **b,** Liver function profile, represented as serum ALT (Left) and AST (Right) in these mice (*n*=5). Error bars represent SD (**a,b**). *P* values were calculated using a two-tailed unpaired Student’s *t*-test (**a,b**).

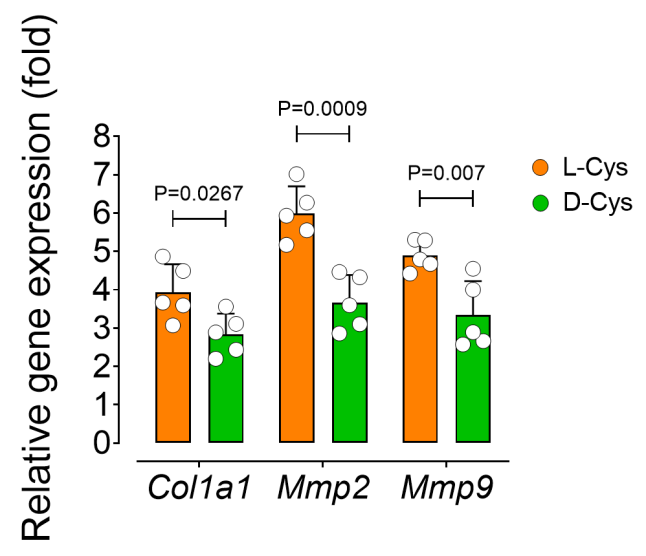

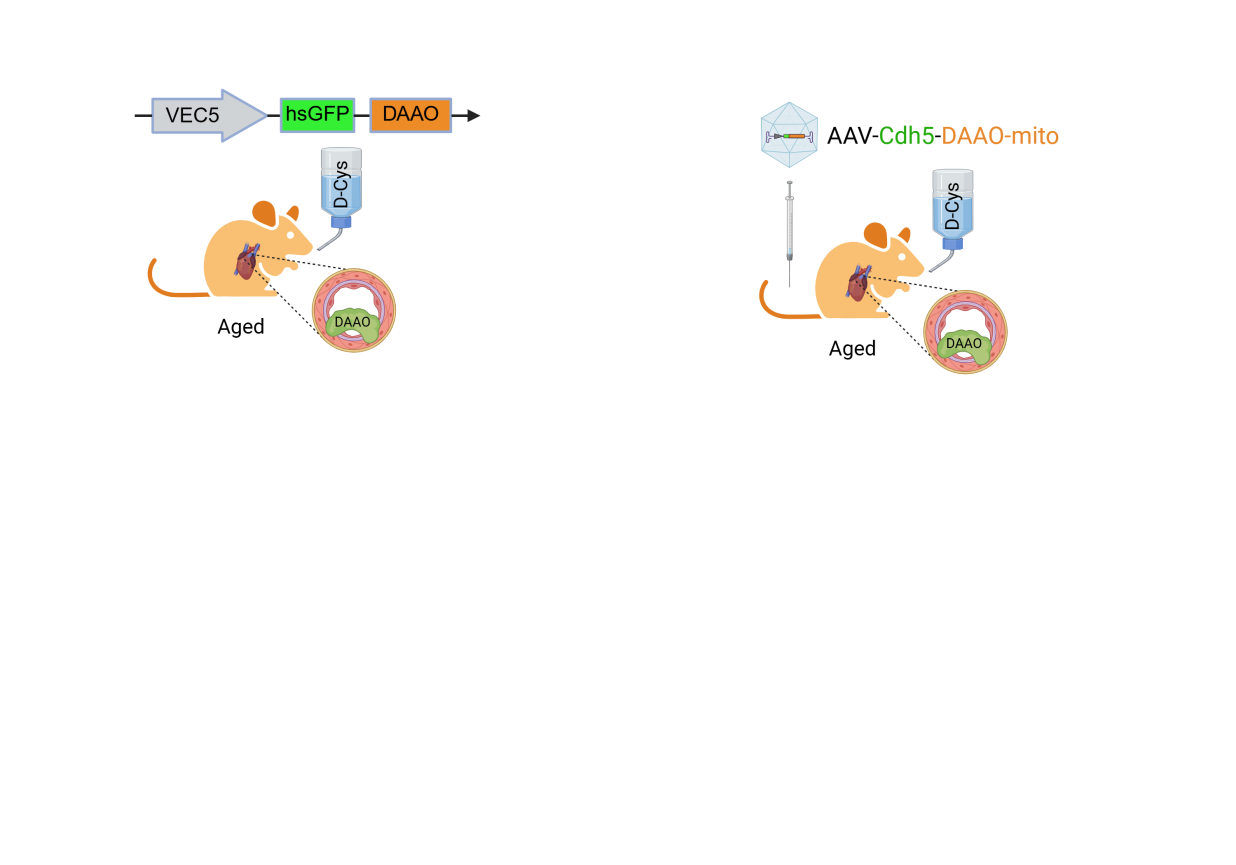

**Supplemental Fig. 5.** **Chemogenetic H_2_S perturbs aortic tissue remodeling in ageing.** qPCR demonstrates transcriptional alterations of the ECM remodeling and pro-atherogenic markers *Mmp2* and *Mmp9* in aortas from aged mice infected with an AAV9-Cdh5-DAAO- mito construct in the presence of 1 M D-Cys or L-Cys for 4 weeks (*n*=5). Error bars represent SD. *P* values were calculated using a two-tailed unpaired Student’s *t*-test.
