## Supplemental Information for "Chemogenetic Hydrogen Sulfide Rescues Vascular Senescence in Ageing"

Supplemental Tables

Supplemental Table 1

Supplemental Table 2

Supplemental Table 3

Supplemental Table 4

Supplemental Table 5

**Supplemental Table 1. Aging Heart Zurich Cohort characteristics**

|  | **Overall cohort (n=187)** | **Control (n=77)** | **ASCVD (n=110)** | **p-value** |
| --- | --- | --- | --- | --- |
| Age (y) | 75.3 ± 11.4 | 63.1 ± 6.4 | 83.8 ± 3.8 | 1.25 × 10^-8^ |
| Sex, F (%) | 60 (33.9%) | 19 (24.7%) | 41 (37.3%) | 0.081 |
| **Cardiovascular risk factors** |  |  |  |  |
| Family history of CV disease, n (%) | 28 (15%) | 7 (9.1%) | 21 (19.1%) | 0.064 |
| Smoking, n (%) | 36 (19.2%) | 4 (5.2%) | 32 (29.1%) | 2.38 × 10^-5^ |
| Alcohol consumption, n (%) | 55 (29.4%) | 5 (6.5%) | 50 (45.4%) | 1.63 × 10^-9^ |
| **Treatments** |  |  |  |  |
| Lipid-lowering therapy, n (%) |  | - | 110 (100%) |  |
| Antihypertensive therapy, n (%) |  | - | 76 (69.1%) |  |
| Anti-platelets & Anticoagulants, n (%) |  | - | 64 (58.2%) |  |
| **Biochemistry** |  |  |  |  |
| Total cholesterol, mmol/L |  | - | 4.12 ± 37.8 |  |
| LDL-C, mmol/L |  | - | 2.4 ± 1.1 |  |
| HDL-C, mmol/L |  | - | 1.4 ± 0.4 |  |
| Triglyceride, mmol/L |  | - | 1.2 ± 0.6 |  |
| Fasting glucose, mg/dL |  | - | 6.8 ± 2.5 |  |
| HbA1c, % |  | - | 5.8 ± 1.1 |  |
| *Troponin T, ng/L |  | - | 83 (39.8–215) |  |
| *NT-ProBNP, pg/mL |  | - | 1488 (613-3387) |  |
| Hemoglobin, g/L |  | - | 111.5 ± 17.5 |  |
| **Inflammatory marker** |  |  |  |  |
| *CRP, mg/L |  | - | 9.5 (1.85-36.5) |  |
| **Biomarkers** |  |  |  |  |
| *H_2_S, µmol/L | 0.198 (0.094-0.399) | 0.332 (0.167-0.472) | 0.121 (0.063–0.205) | <0.0001 |
| **Kidney and Liver function** |  |  |  |  |
| *Creatinine, μmol/L |  | - | 100 (83.25–130.5) |  |
| *eGFR, mL/min/1.73m^2^ |  | - | 54 (43–65.8) |  |
| *ALT, U/L |  | - | 20.5 (14–32) |  |
| *AST, U/L |  | - | 28 (22–45.75) |  |
| *GGT, U/L |  | - | 35.5 (22.25-67.75) |  |
| Data as mean ± SD, n (%) or median and interquartile range (*). P-values were calculated by the two-sided Student t-test (continuous parametric data), two-sided by Mann-Whitney *U*-test (continuous nonparametric data), two-sided Fisher’s Exact test (categorical variables), or two-sided Welch’s t-test. Abbreviations: ALT, alanine aminotransferase; AST, aspartate aminotransferase; CV, cardiovascular disease; F, female; GGT, gamma-glutamyl transferase; GFR, glomerular filtration rate; H_2_S, hydrogen sulfide; HbA1c, glycosylated haemoglobin; HDL-C, high-density lipoprotein cholesterol; CRP, C-reactive protein; LDL-C, low-density lipoprotein cholesterol; NT-ProBNP, B-type natriuretic peptide. | | | | |

**Supplemental Table 2. Validation CAD Study characteristics**

|  | **non-CAD (n=5)** | **CAD (n=5)** |
| --- | --- | --- |
| Age (y) | 61.8 ± 8.3 | 65.6 ± 10.8 |
| Sex, (M/F) | 4/1 | 4/1 |
| **Cardiovascular risk factors** |  |  |
| Family history of CV disease (n) | 0 | 1 |
| Smoking (n) | 0 | 2 |
| **Treatments** |  |  |
| Lipid-lowering therapy (n) | 2 | 5 |
| Antihypertensive therapy (n) | 2 | 2 |
| Anti-platelets & Anticoagulants (n) | 3 | 5 |
| **Biochemistry** |  |  |
| Total cholesterol, mmol/L | 4.4 ± 0.7 | 3.9 ± 0.5 |
| LDL-C, mmol/L | 2.4 ± 0.7 | 1.6 ± 0.9 |
| HDL-C, mmol/L | 2.2 ± 1.6 | 1.5 ± 0.3 |
| Triglyceride, mmol/L | 2.7 ± 2.8 | 1.4 ± 0.4 |
| HbA1c, % | 5.5 ± 0.2 | 5.9 ± 0.7 |
| Troponin T, ng/L | 9.0 ± 5.3 | 58 ± 76.7 |
| NT-ProBNP, pg/mL | 248.8 ± 371.9 | 1241.2 ± 1387 |
| Hemoglobin, g/L | 146.8 ± 10 | 123.8 ± 34 |
| **Inflammatory marker** |  |  |
| CRP, mg/L | 3.4 ± 2.1 | 1.65 ± 1.2 |
| **Biomarkers** |  |  |
| H_2_S, µmol/L | 0.1 ± 0.05 | 0.4 ± 0.1 |
| **Kidney and Liver function** |  |  |
| Creatinine, μmol/L | 111.6 ± 46.2 | 82.6 ± 9.1 |
| eGFR, mL/min/1.73m^2^ | 81.2 ± 9.6 | 75.2 ± 39.2 |
| ALT, U/L | 23.6 ± 10.4 | 45.2 ± 31.5 |
| GGT, U/L | 37.5 ± 28.6 | 50 ± 20 |
| Data as mean ± SD. Abbreviations: ALT, alanine aminotransferase; CV, cardiovascular disease; F, female; GGT, gamma-glutamyl transferase; GFR, glomerular filtration rate; H_2_S, hydrogen sulfide; HbA1c, glycosylated haemoglobin; HDL-C, high-density lipoprotein cholesterol; CRP, C-reactive protein; LDL-C, low-density lipoprotein cholesterol; NT-ProBNP, B-type natriuretic peptide. | | |

**Supplemental Table 3. Western diet composition**

**Western Diet – 0.21% Cholesterol (SSNIFF)**

**HF / High cholesterol diet for mice** | corresponds to TD.88137

**Description:**

The Western diet contains 21% butter fat and approximately 0.21% cholesterol (0.15% supplemented).

This diet will therefore induce hyperlipidemic disorders, hypercholesterolemia, and atherosclerosis—i.e., plaque formation—in *ApoE*- and *Ldlr*-deficient mice.
*The diet is intended for ad libitum feeding. The animals should have free access to fresh water.*

| **Energy Content:**   - **Gross Energy (GE):** 21.8 MJ/kg - **Metabolizable Energy (ME):** 19.1 MJ/kg *(Atwater values; ≈ 4,575 kcal/kg)*   **Energy Distribution:**   - **Fat:** 42 kJ% - **Protein:** 15 kJ% - **Carbohydrates:** 43 kJ%   **Nutritional Composition:**  **Crude Nutrients [%]:**   - Crude protein (N × 6.25): 17.3% - Crude fat: 21.1% - Crude fibre: 5.0% - Crude ash: 4.2% - Starch: 14.4% - Sugar: 34.3% - Nitrogen-free extracts: 49.8%   **Minerals [%]:**   - Calcium: 0.76 - Phosphorus: 0.45 - Ca:P ratio: 1.69:1 - Sodium: 0.24 - Magnesium: 0.10 - Potassium: 0.54   **Amino Acids [%]:**   - Lysine: 1.43 - Methionine: 0.93 - Cystine: 0.07 | - Tryptophan: 0.22 - Met + Cys: 1.00 - Threonine: 0.76 - Arginine: 0.67 - Histidine: 0.52 - Valine: 1.20 - Isoleucine: 0.97 - Leucine: 1.71 - Phenylalanine: 0.89 - Phe + Tyr: 1.80 - Glycine: 0.34 - Glutamic acid: 3.88 - Aspartic acid: 1.28 - Proline: 1.97 - Serine: 1.03 - Alanine: 0.52   **Fatty Acids [%]:**   - C 4:0 (Butyric): 0.80 - C 6:0: 0.53 - C 8:0: 0.29 - C 10:0: 0.63 - C 12:0: 0.72 - C 14:0: 2.22 - C 16:0: 5.60 - C 17:0: 0.14 - C 18:0: 2.05 - C 20:0: 0.04 - C 16:1: 0.38 - C 18:1: 4.65 - C 18:2: 0.38 - C 18:3: 0.11 | **Vitamins (per kg):**   - Vitamin A: 15,000 IU - Vitamin D₃: 1,500 IU - Vitamin E: 150 mg - Vitamin K (as MNB): 20 mg - Thiamine (B₁): 26 mg - Riboflavin (B₂): 16 mg - Pyridoxine (B₆): 16 mg - Cobalamin (B₁₂): 30 μg - Niacin: 49 mg - Pantothenic acid: 55 mg - Folic acid: 16 mg - Biotin: 300 μg - Choline: 920 mg   **Trace Elements (per kg):**   - Iron: 49 mg - Manganese: 22 mg - Zinc: 41 mg - Copper: 11 mg - Iodine: 0.3 mg - Selenium: 0.2 mg   **Cholesterol:**   - 2,070 mg/kg   **Main Products:**  E15721-34: 10 mm pellets  E15721-347: 10 mm pellets, γ-irradiated 25 kGy |
| --- | --- | --- |

**Supplemental Table 4. Nucleotide sequence of primers used in this study**

Gene-specific forward and reverse primers for RT-PCR amplification. *Ccl2*, C-C motif chemokine ligand 2; *Col1a1*, Collagen 1a1; *Gapdh*, Glyceraldehyde 3-phosphate dehydrogenase; *IL1Β*, Interleukin 1-beta; *IL6*, Interleukin 6; *Mmp2*, Matrix metalloproteinase 2; *Mmp9*, Matrix metalloproteinase 9; CDKN2A, p16; *CDKN2D*, p19; *CDKN1A*, p21.

| **Sequence** | **Gene** |
| --- | --- |
| ATCACCAGCAGCAAGTGTC | *Ccl2* |
| AGGTGGTCCATGGAATCCTG |  |
| AAGAACGGAGATGATGGGGAA | *Col1a1* |
| ATCCAAACCACTGAAACCTCT |  |
| AACAGCAACTCCCACTCTTC | *Gapdh* |
| CCTGTTGCTGTAGCCGTATT |  |
| TCCCAAAGTGCTGGGATTAC | *Il1b* |
| GAAAGCTCAGAGAGGAGGAAAG |  |
| GCCTGCATTAGGAGGTCTTT | *Il6* |
| CCTGACACCAGCAAAGGATAA |  |

| **Sequence** | **Gene** |
| --- | --- |
| GCCTCATACACAGCGTCAATCTT | *Mmp2* |
| CGGTTTATTTGGCGGACAGT |  |
| GGAACTCACACGACATCTTCCA | *Mmp9* |
| GAAACTCACACGCCAGAAGAATTT |  |
| GGTTCTCGCAGTACCATTGA | *p16 (CDKN2A)* |
| CTGACTTCTGAGGTGGGTTTAG |  |
| GTCTCTCCGTTTCCCTTTCTTC | *p19 (CDKN2D)* |
| GCTGGGCGCTTTATTTCATTC |  |
| GGGTGCGGTGATGGATAAA | *p21 (CDKN1A)* |
| ACTGCTGAGAACAGGAAGAAC |  |

**Supplemental Table 5. Antibodies and reagents used in this study**

| **Antibodies** | **Reagents and Chemicals** |
| --- | --- |
| anti-p16^INK4a^ (1:100 and 1:1000; abcam, ab211542) | PolyJet transfection reagent (SignaGen Laboratories, SL100688) |
| anti-phospho-Histone H2A.X Ser139 (1:1000; CST, 80312) | EBM-2 culture medium (Lonza, 00190860) |
| anti-IL6 (1:1000; Invitrogen, M-620) | EGM-2 Endothelial Cell Growth Medium-2 BulletKit^TM^ (Lonza, CC-3162/6) |
| anti-CTH (1:1000; CST, 19689) | Trypan Blue (PAN BIOTEC, P08-34100) |
| anti-GFP (1:1000; CST, 2555) | TRIzol Reagent^®^ (Sigma-Aldrich, T9424) |
| anti-Ki67 (1:100; Invitrogen, MA5-14520) | RIPA lysis buffer (ThermoFisher Scientific, 89901) |
| anti-CD31 (1:100; eBioscience, 14-0311-82) | Halt protease-phosphatase inhibitors cocktail (ThermoFisher Scientific, 78442) |
| anti-GAPDH (1:1000; Merck Millipore, G9545) | Bouin’s fixative (Muto Pure Chemicals, 6482.3) |
| conjugated secondary anti-Alexa Fluor^®^ 488 (1:2000; abcam, ab150157) | DAPI (Sigma-Aldrich, D9542) |
| conjugated secondary anti-Alexa Fluor^®^ 594 (1:2000; abcam, ab150080) | Cellular senescence assay kit (Merck Millipore, KAA002) |
| conjugated secondary anti-Alexa Fluor^®^ 647 (1:2000; abcam, ab150075) | BCA protein assay kit (ThermoFisher Scientific, 23225) |
| anti-goat anti-rabbit IgG-HRP (1:2000; Southern Biotechnology, 4030-05) | Cholesterol and triglycerides colorimetric assays (ABX Pentra C400, HORIBA Medical) |
| anti-goat anti-mouse IgG-HRP (1:2000; Southern Biotechnology, 1036-05) | Creatinine colorimetric assay kit (Cayman Chemical, 700460) |
|  | Urea Nitrogen (BUN) colorimetric detection kit (Invitrogen, EIABUN, 15768437) |
